# Isoflurane and surgery aggravate APOE4-dependent lipid dysregulation and neural dysfunction, leading to neurological impairment in male mice

**DOI:** 10.64898/2026.07.31.742028

**Authors:** Yun Li, Yuanyuan Ji, Cosar Uzun, Syed Taufiqul Islam, Meigeng Hu, Dan Zhao, Yaping Li, Hangnoh Lee, Zihui Wang, Hui Li, Jace W. Jones, Shaolin Liu, Junfang Wu

## Abstract

**Purpose:** Perioperative exposure to the volatile anesthetic isoflurane (ISO) has been associated with cognitive and olfactory deficits and may increase the risk of Alzheimer’s disease (AD). Apolipoprotein E4 (APOE4), the strongest genetic risk factor for AD, contributes to disease pathogenesis through disrupted lipid homeostasis. However, whether and how isoflurane interacts with APOE genotype to influence neurological vulnerability remains unclear.

**Methods:** Young adult, presymptomatic humanized APOE4 and APOE3 knock-in mice underwent laparotomy under 2 h of isoflurane anesthesia. Microglia and astrocytes were isolated from the olfactory bulb (OB) and hippocampus (HI) by magnetic-activated cell sorting. Lipid composition, transcriptional responses, and functional outcomes were assessed using lipidomic, bulk RNA-seq, and longitudinal behavioral testing. *In vivo* and *ex vivo* electrophysiological recordings evaluated neuronal excitability and synaptic transmission in both regions.

**Results:** By day 7 post-anesthesia, cell type-specific lipidomic profiling of both OB and HI revealed more pronounced lipid perturbations in microglia and astrocytes from APOE4/ISO mice than from APOE3 mice, characterized by elevated free fatty acids, increased lipid peroxidation, triglyceride depletion, and reduced hippocampal hexosylceramides and cardiolipins. Electrophysiological recordings showed greater olfactory circuit dysfunction in APOE4/ISO mice, accompanied by persistent odor memory deficits, transient olfactory sensitivity loss, early motor coordination impairments, and delayed cognitive deficits. RNA sequencing of the OB identified downregulated lipid metabolism and atherosclerosis-related pathways.

**Conclusion:** These findings establish a mechanistic link between APOE4-dependent glial lipid dysregulation, olfactory circuit dysfunction, and delayed cognitive impairment following isoflurane anesthesia and surgery, highlighting lipid homeostasis as a potential therapeutic target.

## 1 Introduction

General anesthesia (GA) is essential for modern surgery, but perioperative exposure to volatile anesthetics and surgical stress may contribute to postoperative neurocognitive dysfunction (PND), characterized by impairments in memory, attention, and executive function [1–3]. Anesthesia and surgery can also induce olfactory dysfunction (OD), and emerging evidence suggests that postoperative OD may precede or associate with later cognitive decline [4–7]. Beyond its impact on quality of life, OD is increasingly recognized as an early marker of neurodegenerative disease, including Alzheimer’s disease (AD) [8–10]. Longitudinal studies indicate that OD can precede cognitive decline by years or decades [11, 12], potentially through olfactory bulb (OB)-linked propagation of pathogenic proteins and disrupted connectivity with limbic memory circuits [6, 10].

Preclinical studies support anesthesia-associated neurotoxicity, including Tau phosphorylation after repeated sevoflurane exposure and long-term memory impairment after isoflurane (ISO) exposure in rodents [13–16]. However, how transient perioperative insults might interact with genetic risk to promote later AD-related vulnerability remains unclear. The apolipoprotein E4 (APOE4, E4) is the strongest common genetic risk factor for sporadic AD, conferring an approximately threefold increased risk in heterozygotes and a 10- to 15-fold increased risk in homozygotes compared with non-carriers [17, 18]. Approximately 25-30% of U.S. adults carry at least one APOE4 allele [19, 20]. Clinical and preclinical studies suggest that APOE4 may increase susceptibility to PND, but direct apolipoprotein E3 (APOE3, E3) versus APOE4 comparisons remain limited [21–25]. Humanized E4 knock-in (KI) mice recapitulates key aspects of E4-driven vulnerability without overexpressing mutant amyloid or tau, providing a genetically precise and translationally relevant platform for studying anesthesia-related neurotoxicity [26, 27]. APOE4 is linked to age-dependent changes in glial lipid metabolism, synaptic integrity, and network excitability, including lipid droplet accumulation and impaired glial support of neuronal function [28, 29]. Yet glial lipid dysregulation remains underexplored in PND models, and whether perioperative exposure interacts with APOE genotype to shape postoperative neurological risk is unknown.

Therefore, we hypothesized that APOE4 interacts with isoflurane exposure and surgical stress to exacerbate neurological disruption. To test this, presymptomatic young adult APOE3 and APOE4 knock-in mice underwent laparotomy with 2 h of isoflurane anesthesia. Using integrated lipidomic, electrophysiological, behavioral, and transcriptomic analyses, we identified APOE4-specific vulnerabilities marked by early glial lipid dysregulation, olfactory circuit dysfunction, and persistent sensory deficits that preceded later cognitive impairment. These findings establish olfactory dysfunction as an early, accessible marker of E4 susceptibility, highlighting a window for intervention to mitigate anesthesia- and age-related cognitive decline.

## 2 Methods

### 2.1 Mouse general anesthesia and operation

All experimental procedures were approved by the Institutional Animal Care and Use Committees of the University of Maryland School of Medicine and the University of Georgia. Homozygous humanized APOE4 KI [B6(SJL)-Apoetm1.1(^APOE*4^)Adiuj/J, stock no. 027894] and APOE3 KI [B6.Cg-Apoeem2^(APOE*)^Adiuj/J, stock no. 029018] mice (Mus musculus; male; 10-14 weeks old; 20-25 g) were obtained from The Jackson Laboratory. Mice were subjected to a perioperative paradigm consisting of 2 h ISO anesthesia with laparotomy operation (OP) to model surgical stress, as detailed in supplemental materials. Sham controls remained in their home cages and were exposed to room air for 2 h, to better model clinically relevant baseline conditions.

### 2.2 Adult cell isolation

At day 7 after ISO/OP, mice were perfused with 50 ml ice-cold saline. OB and hippocampus (HI) tissue from three mice of the same genotype and treatment was pooled per biological replicate. CD11b⁺ microglia and ACSA2⁺ astrocytes were isolated by magnetic-activated cell sorting according to the manufacturer’s protocol (Miltenyi Biotec).

### 2.3 Lipidomics

Lipidomic analysis of astrocytes and microglia samples was performed as described in previous publications [30, 31]. Data acquisition was performed using Thermo Xcalibur software, and data were processed with Xcalibur 4.2 and TraceFinder 5.1. Additional statistical and visualization analyses were carried out using Prism 10 (GraphPad) and MetaboAnalyst 6.0.

### 2.4 *In vivo* electrophysiological recording and data analysis

Surgical preparations for *in vivo* electrophysiological recordings and data analysis were performed as previously described with modifications [7, 32, 33]. To assess the short-term and long-term effects of ISO and surgery, recordings were performed 24 hours or 7 days after the GA cessation, respectively. Spontaneous neuronal excitability and network activities were recorded as described previously [32, 33], but with a 32-channel neural probe (60 μm shank width, 15 μm shank thickness and 50 μm inter-channel distance) inserted into the MCL on the medial side of each OB. Data from *in vivo* electrophysiology experiments are presented as mean ± SEM. Statistical analysis was performed using the Origin 2024 software. Comparison of cumulative probability distribution between two groups was analyzed using the nonparametric Mann-Whitney U-test. Spikes were sorted from the raw data with Offline Sorter V4 software (Plexon). The separation of different units was performed by principal component analysis based on criteria of waveform parameters and multidimensional clusters. Both spike and LFP data were further analyzed with NeuroExplorer V5.204 (Nex Technologies) and graphed with Excel and Origin 2024 (OriginLab Corporation, Northampton, MA). Instantaneous spike frequency (ISF) calculated from inter-spike interval of individual units was utilized to assess neuronal excitability. At the population level, an average of ISF across multiple units was used to present each recording. A cumulative probability of ISF averaged across multiple animals in each group was applied to compare neuronal excitability among multiple groups. Power spectrum density (PSD) of LFP signals ranging from 0.1 to 200 Hz, which covers different frequency bands including theta (0-12 Hz), beta (12-30 Hz), and gamma (low gamma, 30-60; high gamma 60-100 Hz), was measured to reflect the strength of oscillatory activities that are derived from interactions among excitatory and inhibitory neurons thus reflect network operation [33–35]

### 2.5 Patch clamp recording in brain slices and data analysis

Acute OB or brain slices containing HI (350 μm thick) were prepared from E3 or E4 mice as previously described [36, 37]. Cumulative probability distribution plots of sEPSCs or sIPSCs were created in Origin 2024. Comparison between two groups was analyzed using the non-parametric Kolmogorov-Smirnov test. Three-way ANOVA was used to analyze membrane resistance, resting membrane potential, and current injection-evoked spikes. p ˂ 0.05 was considered statistically significant. Patch pipettes (5-7 MΩ) were pulled from thin-walled glass capillaries with filament (Sutter Instrument, Novato, CA). The internal solution contained (in mM): 117 K-gluconate, 10 KCl, 10 HEPES, 4 EGTA, 12 KOH, 0.5 CaCl₂, 3 Mg-ATP, 0.3 Na₂-GTP, and 7 Na₂-phosphocreatine (pH adjusted to 7.26, 290 mOsm). Automatic detection of spontaneous excitatory or inhibitory postsynaptic currents (sEPSCs or sIPSCs) was performed as previously described with Wdetecta (https://hlab.stanford.edu/wdetecta.php) followed by manual checks to ensure accuracy [36]. Instantaneous sEPSC or sIPSC frequencies were then calculated from their time stamp data generated from Wdetecta detection.

### 2.6 Assessment of neurological function

Behaviour was assessed longitudinally for 88 days by investigators blinded to genotype. Treatment allocation could not be blinded because of the visible surgical incision (Supplementary Figure S1). Groups compared include E3/Sham, E4/Sham, E3/ISO, E4/ISO, with units being a single mouse. Detailed methods and specific tests can be found in supplementary materials.

### 2.7 RNA extraction and bulk RNA sequencing (RNAseq)

At 90 days after ISO/OP, mice were perfused with 50 ml ice-cold saline, and total RNA was isolated from OB and HI using the miRNeasy kit (Qiagen, Cat. No. 217084). Poly(A)-enriched libraries were prepared by Novogene and sequenced as 150-bp paired-end reads on an Illumina NovaSeq 6000.

### 2.8 Statistical analysis

Data are reported as mean (SEM), with individual values shown where applicable. Normality was assessed using the Shapiro-Wilk test. Electrophysiological and behavioral data were analyzed using ANOVA-based approaches, whereas molecular, transcriptomic, and lipidomic datasets used assay-specific methods. Statistical models, post hoc tests, multiple-testing corrections, sample sizes, and significance thresholds are reported in the relevant Methods subsections and figure legends. Additional design and inclusion details are provided in the Supplemental Information. Unless otherwise specified, P<0.05 was considered significant.

Full details on methods are provided in the Supplementary data.

## 3 Results

### 3.1 APOE4 microglia and astrocytes show heightened lipid disruption post-ISO/OP

To investigate the effects of ISO/OP on lipid composition, young adult E4 and E3 mice were exposed to 2 h of ISO/OP followed by a 7-day recovery period. The OB and HI were subsequently dissected, and microglia and astrocytes were isolated for lipidomic analysis. This approach is particularly relevant because the effects of E4 on lipid metabolism are thought to be mediated primarily through glial cells, where it disrupts membrane composition, lipid trafficking, and inflammatory signaling. We first examined the influence of genotypes by directly comparing E4 with E3 mice following ISO/OP exposure. In OB astrocytes, PCA and differential analysis revealed genotype-dependent alterations in phosphatidylcholines (PC), phosphatidylethanolamine (PE), and Cer species (Fig. 1A-C, n=3 samples/9 mice/group, each sample was pooled from three mice). OB microglia showed similarly robust differences, with broad shifts across multiple lipid classes and subtypes (Fig. 1D-F). Although HI astrocytes demonstrated partial overlap between E3 and E4 groups, changes in sphingomyelin (SM) and PE remained more pronounced than those observed in bulk tissue (Fig. 1G-I). In contrast, HI microglia displayed clear separation between E4/ISO and E3/ISO, with elevated PE and SM emerging as distinctive features of the E4 lipid profile (Fig. 1J-L).

**Fig. 1.**
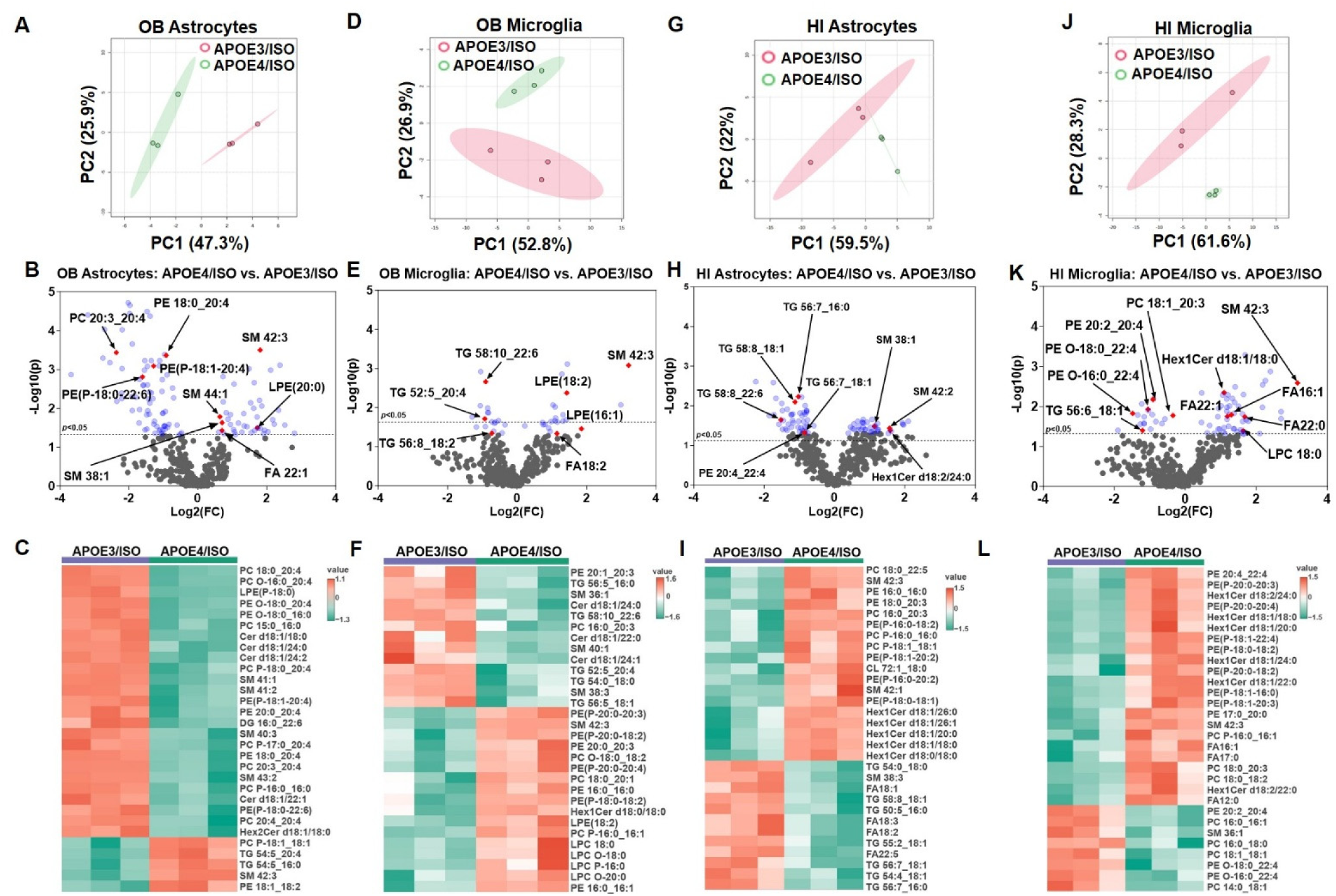
APOE4 drives heightened lipid dysregulation in olfactory bulb (OB) and hippocampal (HI) astrocytes and microglia 7 days after isoflurane (ISO) exposure and laparotomy operation (OP). **A-C** OB astrocytes, **D-F** OB microglia, **G-I** HI astrocytes, and **J-L** HI microglia from APOE4/ISO vs. APOE3/ISO mice at 7 days post-exposure. In each set, PCA illustrates group separation, with volcano plots and heatmaps depicting differentially abundant lipids. n=3 samples/9 mice/group. Each sample was pooled from three male mice, with a total of nine mice per group.

We next sought to isolate the effects of ISO/OP in a specific genotype. In the OB (Supplementary Figure S2A-F), PCA revealed clear separation between E4/ISO and E4/Sham groups in both astrocytes and microglia. Differential expression analysis highlighted cell type-specific differences: astrocytes displayed increases in SM, lysophosphatidylcholine (LPC) and lysophosphatidylethanolamine (LPE) species, accompanied by a pronounced reduction in PE, whereas microglia showed reduced PE, with concomitant upregulation of SM and LPE species. In the HI, astrocytes and microglia exhibited even stronger treatment-based clustering than bulk assays (Supplementary Figure S2G-L). Astrocytes showed decreased PE, while also displaying marked elevations in SM, FA, LPC and LPE. Similarly, microglia also showed an overall decrease of PE lipids, while FA, LPC and SM types were elevated after ISO/OP. Comparable disruptions were also detected in E3 astrocytes and microglia at 7 d after ISO/OP (Supplementary Figure S3), underscoring the broad impact of anesthesia on glial lipid metabolism. While baseline differences between E3 and E4 were detectable (Supplementary Figure S4), the combined analysis of lipid classes and subtypes pointed to a strong interaction between genotype and ISO/OP exposure as a driver of glial lipid dysregulation. In summary, ISO/OP drives region- and genotype-dependent disruptions to lipid metabolism in both astrocytes and microglia, with E4 glia showing more pronounced alterations than glia from E3. These distinct cellular signatures underscore glial lipid metabolism as a key node of vulnerability and set the stage for linking such changes to functional outcomes in later analyses.

### 3.2 APOE4 impairs neuronal excitability in the OB of awake mice after ISO exposure

Lipids modulate ion channels, receptors, and neuronal excitability [38, 39]. Next, we asked whether ISO-associated glial lipid alterations were accompanied by changes in OB excitability. We performed *in vivo* recordings from the mitral cell layer using 32-channel linear probes in awake, head-fixed mice on days 1 and 7 after 2 h ISO exposure (Supplementary Figure S5). Spontaneous spiking and LFP oscillatory activity were recorded across channels to assess neuronal excitability and network function.

On day 1 after exposure (Fig. 2A-B), ISO increased mitral cell spiking frequency in E3 mice compared with sham controls (n=6 recordings from 3 ISO mice; n=7 recordings from 3 sham mice), indicating hyperexcitability. In contrast, ISO reduced spiking frequency in E4 mice compared with E4 sham controls (n=6 recordings from 3 mice/group), with no baseline difference between E3 and E4 sham groups. Thus, ISO produced opposing genotype-dependent effects on mitral cell excitability on day 1. This divergent pattern persisted on day 7 (Fig. 2C-D). ISO increased spiking frequency in E3 mice compared with E3 sham controls (n=5 recordings from 3 ISO mice; n=8 recordings from 3 sham mice), but reduced spiking frequency in E4 mice compared with E4 sham controls (n=8 recordings from 3 ISO mice; n=6 recordings from 3 sham mice). Although E3 and E4 sham groups also differed on day 7, ISO-exposed E4 mice still showed significantly lower spiking frequency than ISO-exposed E3 mice, supporting a persistent genotype-specific response to ISO.

**Fig. 2.**
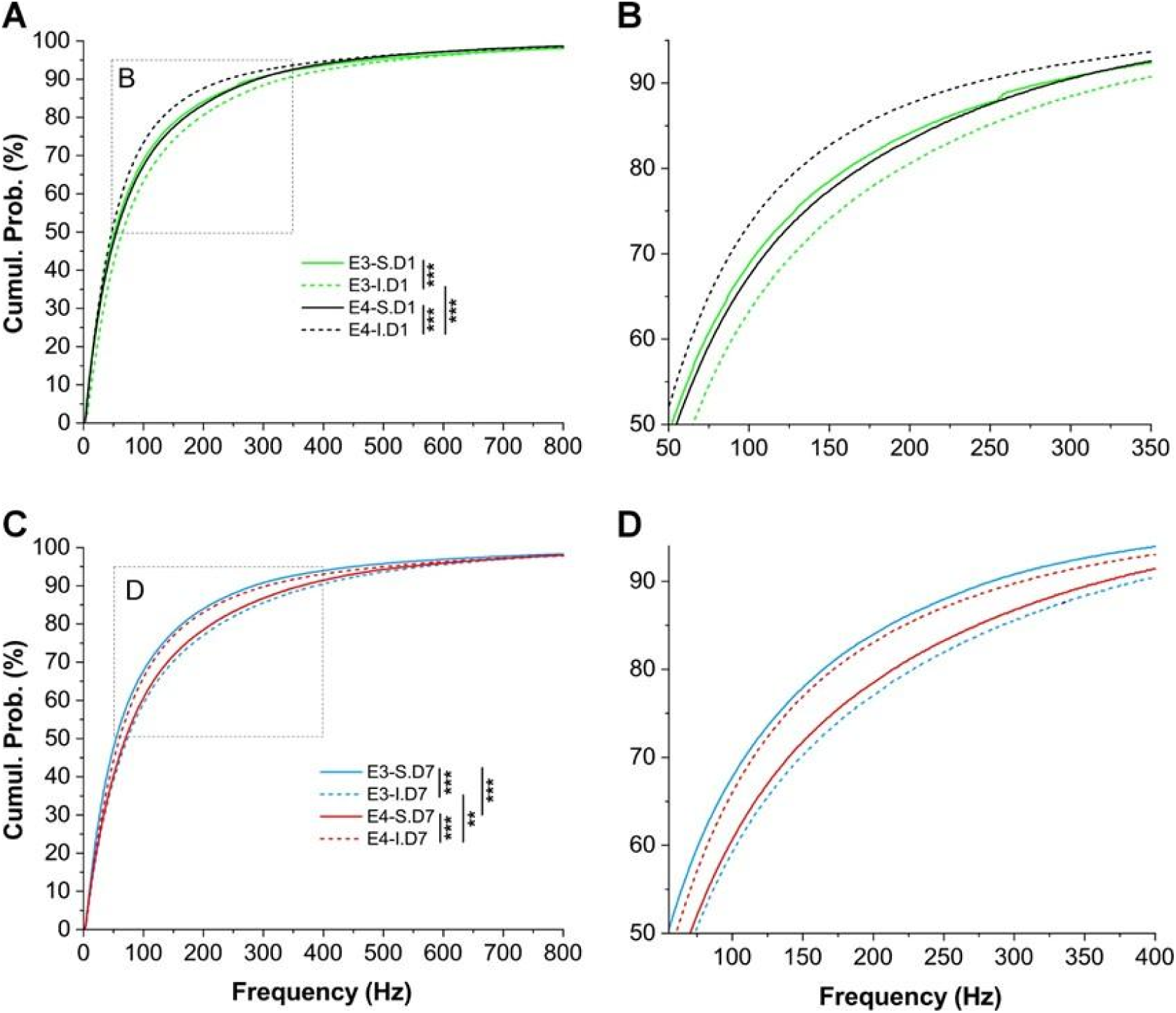
Isoflurane (ISO) effects on neuronal excitability in the OB mitral cell layer on post-ISO day 1 and 7. **A** Comparison of the cumulative probability of spiking frequency on day 1 (D1) post-ISO exposure among four groups of mice: (1) E3/sham (E3-S.D1, solid green curve, n=7 recordings in 3 mice), (2) E3/ISO (E3-I.D1, dashed green curve, n=6 recordings in 3 mice), (3) E4/sham (E4-S.D1, solid black curve, n=6 recordings in 3 mice), (4) E4/ISO (E4-I.D1, dashed black curve, n=6 recordings in 3 mice). **B** Blown-up from the dashed box in A. **C** Comparison of the cumulative probability of spiking frequency on day 7 (D7) post-ISO exposure among four groups of mice: (1) E3/sham (E3-S.D7, solid blue curve, n=8 recordings in 3 mice), (2) E3/ISO (E3-I.D7, dashed blue curve, n=5 recordings in 3 mice), (3) E4/sham (E4-S.D7, solid red curve, n=6 recordings in 3 mice), (4) E4/ISO (E4-I.D7, dashed red curve, n=8 recordings in 3 mice). **D** Blown-up from the dashed box in C. Comparison between the two groups was analyzed using the Mann-Whitney test. **p<0.001, ***p<0.0001.

To determine whether ISO effects on neuronal excitability changed over time, we compared spiking frequency between post-ISO days 1 and 7 within each genotype (Supplementary Figure S6). Time-course analysis showed no difference between E3 sham groups, whereas ISO-exposed E3 mice showed higher spiking frequency on day 7 than day 1, suggesting that ISO-associated hyperexcitability increased over time in E3 mice. In E4 mice, both sham and ISO-exposed groups showed higher spiking frequency on day 7 than day 1, making it difficult to separate ISO effects from time-dependent changes. Together, these findings indicate that a single 2 h ISO exposure produces persistent but divergent effects on OB mitral cell excitability, increasing activity in E3 mice while suppressing activity in E4 mice.

### 3.3 APOE4 exacerbates ISO-induced neuronal network dysfunction in the OB of awake mice

We next analyzed OB local field potentials recorded with 32-channel probes from awake, head-fixed mice to determine whether ISO altered network oscillations. On post-ISO day 1, E3 mice showed increased oscillatory power in the theta (5.5-8.5 Hz, p<0.004) and beta (12-25 Hz, p<0.008) ranges compared with sham controls, with no gamma-band difference (Fig. 3A; n=6 recordings from 3 ISO mice, n=7 recordings from 3 sham mice). In contrast, E4 mice showed reduced PSD at 0-6.5 Hz (p<0.01) and 12-30 Hz (p<0.004), but increased PSD at 46-57 Hz (p<0.004) and 63-100 Hz (p<0.0001) compared with E4 sham controls (n=6 recordings from 3 mice/group). E4 sham mice also showed higher baseline PSD than E3 sham mice across much of the 0-100 Hz range, and ISO-exposed E4 mice showed stronger 28-100 Hz activity than ISO-exposed E3 mice. By post-ISO day 7, genotype-dependent effects persisted. E3/ISO mice showed increased PSD in the 5.5-8 Hz (p<0.02) and 72-80 Hz (p<0.0004) ranges compared with E3 sham controls (Fig. 3B; n=5 ISO recordings from 3 mice, n=8 sham recordings from 3 mice). E4/ISO mice showed broader network disruption, with elevated gamma-band power from 45-100 Hz and reduced beta power from 16-26 Hz compared with E4 sham controls (n=8 ISO recordings from 3 mice, n=6 sham recordings from 3 mice). Baseline E3-E4 sham differences were minimal at day 7, except for a modest increase at 5.5-7.5 Hz in E4 mice (p<0.02), suggesting that the persistent ISO-associated changes in E4 mice were not solely attributable to genotype. Direct comparison of ISO-exposed groups further showed stronger gamma activity from 30-100 Hz in E4 mice (p<0.0001), with weaker 5.5-8 Hz power compared with E3 mice (p<0.02).

**Fig. 3.**
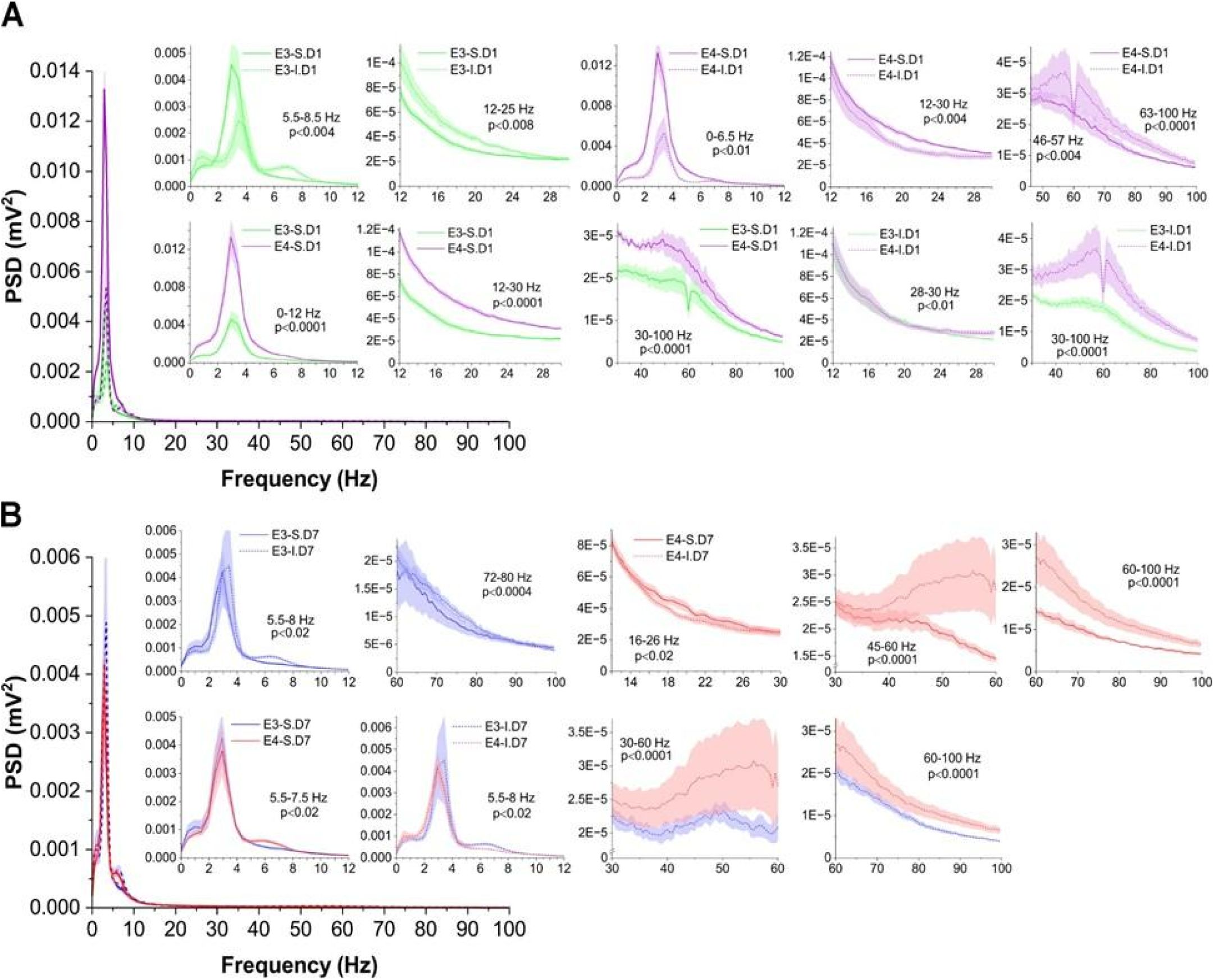
Isoflurane (ISO) effects on neural oscillations of local field potentials on post-treatment day 1 and 7. **A** Comparison of the power spectral density (PSD) of LFP oscillatory activities on day 1 (D1) post-ISO-exposure among four groups of mice: (1) E3/sham (E3-S.D7, solid green curve, n=7 recordings in 3 mice), (2) E3/ISO (E3-I.D7, dashed green curve, n=6 recordings in 3 mice), (3) E4/sham (E4-S.D7, solid purple curve, n=6 recordings in 3 mice), (4) E4/ISO (E4-I.D7, dashed purple curve, n=6 recordings in 3 mice). **B** Comparison of the power spectral density (PSD) of LFP oscillatory activities on day 7 (D7) post-ISO-exposure among four groups of mice: (1) E3/sham (E3-S.D7, solid blue curve, n=8 recordings in 3 mice), (2) E3/ISO (E3-I. D7, dashed blue curve, n=5 recordings in 3 mice), (3) E4/sham (E4-S. D7, solid red curve, n=6 recordings in 3 mice), (4) E4/ISO (E4-I.D7, dashed red curve, n=8 recordings in 3 mice). Insets are zoom-in from A or B highlighting comparisons within difference frequency bands δ/θ/α (a, 0-12 Hz), β (b, 12-30 Hz), low gamma, (c, 30-60 Hz), and high gamma (d, 60-100 Hz). Each curve (solid or dotted) with shaded area presents mean ± SE. Statistical analysis and comparison were conducted using the Mann-Whitney U test.

Time-course analysis showed that ISO-exposed E3 mice had increased 40-60 Hz PSD on day 7 compared with day 1 (Supplementary Figure S7), whereas ISO-exposed E4 mice showed reduced PSD at 12-15 Hz (p<0.04) and 70-100 Hz (p<0.0001) over the same interval. Together, these findings indicate that a single 2 h ISO exposure produces genotype-dependent OB network disruption, with broader and more persistent oscillatory abnormalities in E4 mice that may contribute to olfactory behavioral deficits.

### 3.4 ISO alters synaptic transmission in the OB and HI in E4 mice

Given that ISO can modulate synaptic transmission [40], and that we observed lipid and OB network alterations after ISO exposure, we tested whether ISO altered excitatory and inhibitory inputs onto OB external tufted cells and hippocampal CA1 pyramidal neurons on day 7 after exposure (Figs. 4-5; Supplementary Figure S8-9). Whole-cell recordings measured spontaneous excitatory and inhibitory postsynaptic currents at −60 mV and 0 mV, respectively, with DNQX and gabazine confirming AMPA- and GABA_A_-mediated currents. ETC membrane resistance was lower in E4 sham mice than in E3 sham or E4/ISO mice in the OB, whereas no corresponding differences were detected in HI PNs (Supplementary Figure S8-9).

**Fig. 4.**
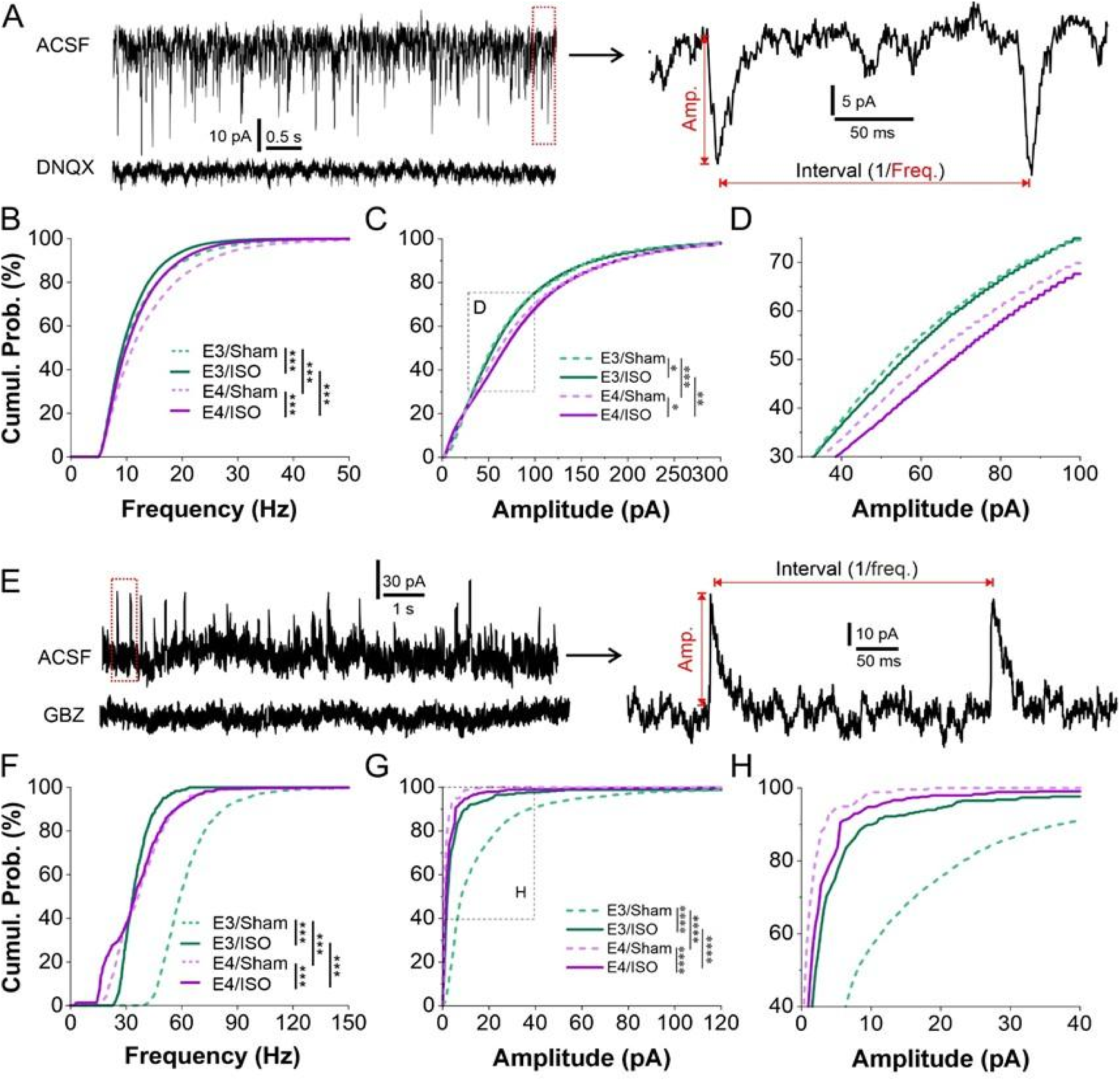
Effects on spontaneous excitatory (sEPSCs) and inhibitory (sIPSCs) postsynaptic currents in external tufted cells (ETCs) in the olfactory bulb (OB). **A** left: typical voltage clamp recording traces showing sEPSCs (top) that were eliminated by bath application of 10 mM DNQX, a selective AMPA receptor blocker (bottom); right: blown-up from left illustrating the measurement of sEPSC amplitude (amp.) and inter-EPSC interval. **B-C** comparisons of sEPSC frequency (B) and amplitude (C) among four groups of mice: E3/Sham (n=8 cells in 2 mice), E3/ISO n=5 cells in 3 mice), E4/Sham (n=9 cells in 3 mice), E4/ISO (n=5 cells in 3 mice). **D** Blown-up from the dashed box area in C. **E** left: typical voltage clamp recording traces showing sIPSCs (top) that were eliminated by bath application of 10 mM gabazine (GBZ), a selective GABA_A_ receptor blocker (bottom); right: blown-up from left illustrating the measurement of sIPSC amplitude (amp.) and inter-IPSC interval. **(F-G)** comparisons of sIPSC frequency (F) and amplitude (G) among four groups of mice: E3/Sham (n=5 cells in 2 mice), E3/ISO (n=4 cells in 3 mice), E4/Sham (n=5 cells in 3 mice), E4/ISO (n=7 cells in 3 mice). **(H)** Blown-up from the dashed box area in G. Comparison between two groups was analyzed using Kolmogorov-Smirnov (K-S) test. *p<0.05, **p<0.01, ***p<0.001, ****p<0.0001.

**Fig. 5.**
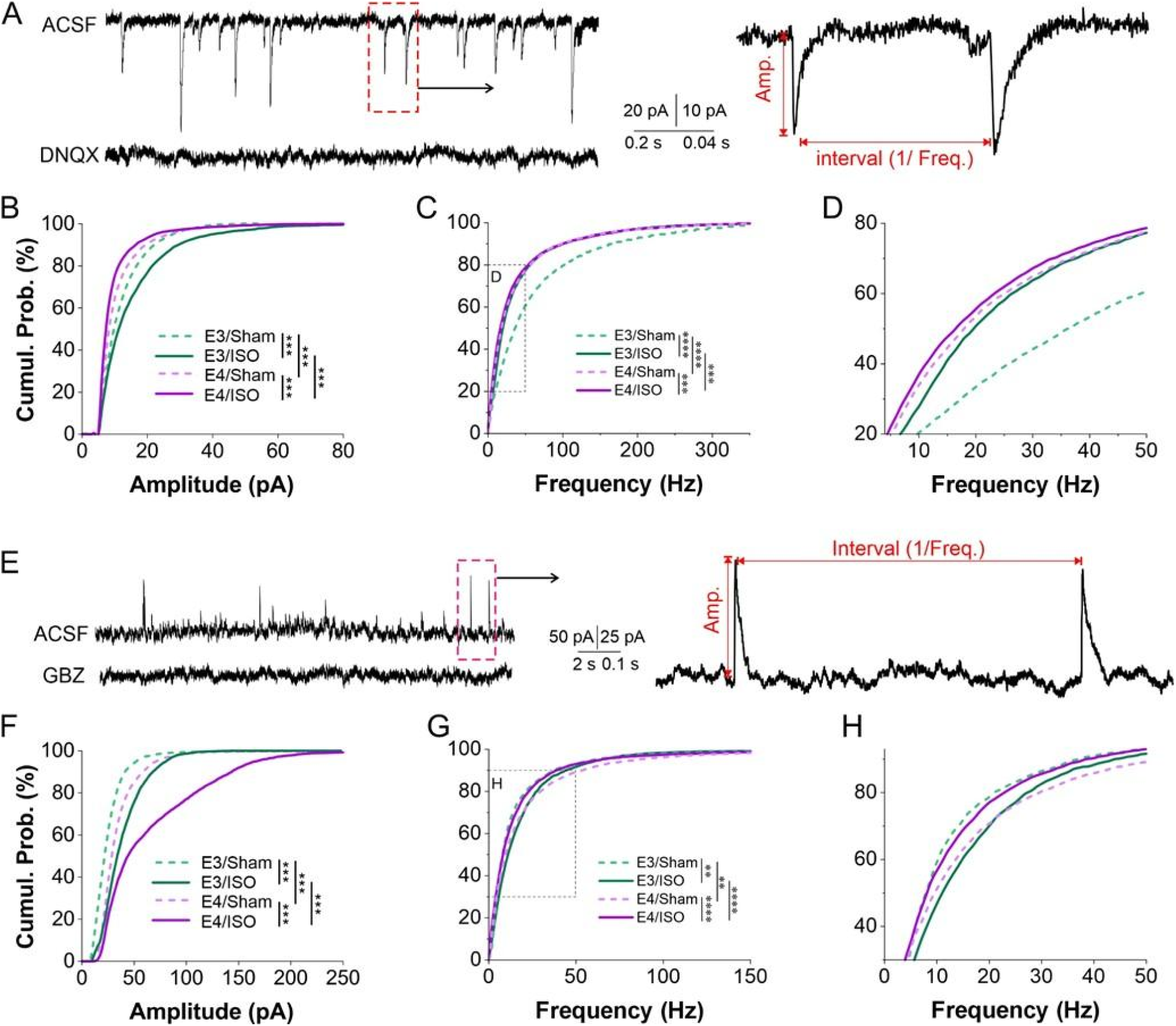
Effects on spontaneous excitatory (sEPSCs) and inhibitory (sIPSCs) postsynaptic currents in pyramidal neurons in the hippocampus (HI). **A** left: typical voltage clamp recording traces showing sEPSCs (top) that were eliminated by bath application of 10 mM DNQX, a selective AMPA receptor blocker (bottom); right: blown-up from left illustrating the measurement of sEPSC amplitude (amp.) and inter-EPSC interval. **B-C** comparisons of sEPSC amplitude (B) and frequency (C) among four groups of mice: E3/Sham (n=8 cells in 2 mice), E3/ISO (n=9 cells in 3 mice), E4/Sham (n=2 cells in 3 mice), E4/ISO (n=15 cells in 3 mice). **D** Blown-up from the dashed box area in C. **E** left: typical voltage clamp recording traces showing sIPSCs (top) that were eliminated by bath application of 10 mM gabazine (GBZ), a selective GABA_A_ receptor blocker (bottom); right: blown-up from left illustrating the measurement of sIPSC amplitude (amp.) and inter-IPSC interval. **F-G** comparisons of sIPSC amplitude (F) and frequency (G) among four groups of mice: E3/Sham (n = 8 cells in 2 mice), E3/ISO (n=7 cells in 3 mice), E4/Sham (n=12 cells in 3 mice), E4/ISO (n = 12 cells in 3 mice). **(H)** Blown-up from the dashed box area in G. Comparison between two groups was analyzed using Kolmogorov-Smirnov (K-S) test. *p<0.05, **p<0.01, ***p<0.001, ****p<0.0001.

At the synaptic level of the OB (Fig. 4), spontaneous excitatory and inhibitory postsynaptic currents were recorded from ETCs and PNs voltage-clamped at −60 mV or 0 mV, respectively (Fig. 4A). Complete blockades by DNQX or gabazine confirmed that sEPSCs and sIPSCs were mediated by AMPA and GABA_A_ receptors, respectively. In OB ETCs, E4 sham mice showed significantly increased sEPSC frequency and amplitude compared with E3 sham mice (Fig. 4B-D), suggesting enhanced synaptic excitation. ISO exposure reduced sEPSC frequency in both E3 and E4 mice, with modest increases in sEPSC amplitude, indicating an overall reduction in synaptic excitation. Similar to sham mice, E4/ISO mice showed higher sEPSC frequency and amplitude than E3/ISO mice, consistent with elevated synaptic excitation in E4 mice. In contrast, E4/Sham mice showed significantly reduced sIPSC frequency and amplitude compared with E3/Sham mice (Fig. 4E-H), suggesting reduced synaptic inhibition. ISO exposure reduced both sIPSC frequency and amplitude in E3 mice, whereas in E4 mice ISO reduced sIPSC frequency but increased sIPSC amplitude, suggesting opposing effects on inhibitory input strength and event frequency in OB ETCs.

Hippocampal CA1 PNs showed a distinct pattern (Fig. 5). E4 sham mice had significantly reduced sEPSC amplitude and frequency compared with E3 sham mice (Fig. 5A-D), indicating reduced hippocampal synaptic excitation. ISO exposure increased sEPSC amplitude in E3 mice but reduced it in E4 mice, indicating opposing genotype-dependent effects. Similar to OB ETCs, ISO reduced sEPSC frequency in hippocampal PNs of both E3 and E4 mice. Among ISO-exposed groups, E4 mice showed lower sEPSC amplitude and frequency than E3 mice, consistent with reduced synaptic excitation. For inhibitory transmission, E4 sham mice showed increased sIPSC amplitude and frequency compared with E3 sham mice (Fig. 5E-H), opposite to the pattern observed in OB ETCs. ISO increased both sIPSC frequency and amplitude in E3 mice, whereas in E4 mice it increased sIPSC amplitude but reduced sIPSC frequency. Among ISO-exposed groups, E4 mice showed higher sIPSC amplitude but lower sIPSC frequency than E3 mice, again differing from the OB ETC pattern.

In summary, E4 increased synaptic excitation and reduced inhibition in OB ETCs, shifting the balance toward hyperexcitation. In contrast, ISO reduced both excitation and inhibition in E3 mice and produced mixed effects in E4 mice. In hippocampal CA1 PNs, E4 had the opposite baseline effect, reducing excitation and increasing inhibition. ISO shifted hippocampal synaptic balance toward inhibition in both APOE genotypes, while affecting excitation differently: increasing sEPSC amplitude in E3 mice but reducing it in E4 mice.

### 3.5 APOE4/ISO mice show persistent olfactory and cognitive deficits linked to chronic transcriptomic dysregulation

Finally, we assessed the long-term neurological effects of ISO/OP in asymptomatic young adult mice. Three-month-old male E3 and E4 mice underwent abdominal surgery followed by 2 h ISO exposure or sham treatment, and behavioral outcomes were measured longitudinally for up to 12 weeks after the intervention. Olfactory memory was assessed by the odor memory (OM) test, in which reduced exploration of a reintroduced cinnamon odor indicates odor recognition. E4/ISO mice showed persistent olfactory memory impairment beginning on day 11 and lasting for several weeks after ISO/OP (Fig. 6A). Buried food testing further showed reduced olfactory sensitivity in E4/ISO mice compared with sham controls and E3 littermates, although this deficit was transient and limited to days 12 and 37 (Fig. 6B). Rotarod testing revealed early locomotor deficits in E4/ISO mice during the first 2 weeks, which were resolved thereafter (Fig. 6C), while nest-building, grip strength, and open-field testing showed no group differences (Supplementary Figure S10-11). Cognitive testing showed no early Y-maze differences, but E4/ISO mice had reduced spontaneous alternation, and increased arm returns on day 86, without changes in total arm entries (Fig. 6D-F). NOR performance did not differ overall, although some E4/ISO mice showed evidence of declarative memory deficits at day 88, and transient impairment was detected during the choice phase on day 41 (Supplementary Figure S10, 12). These findings suggest that ISO/OP induces persistent olfactory deficits and delayed cognitive vulnerability in young adult E4 mice.

**Fig. 6.**
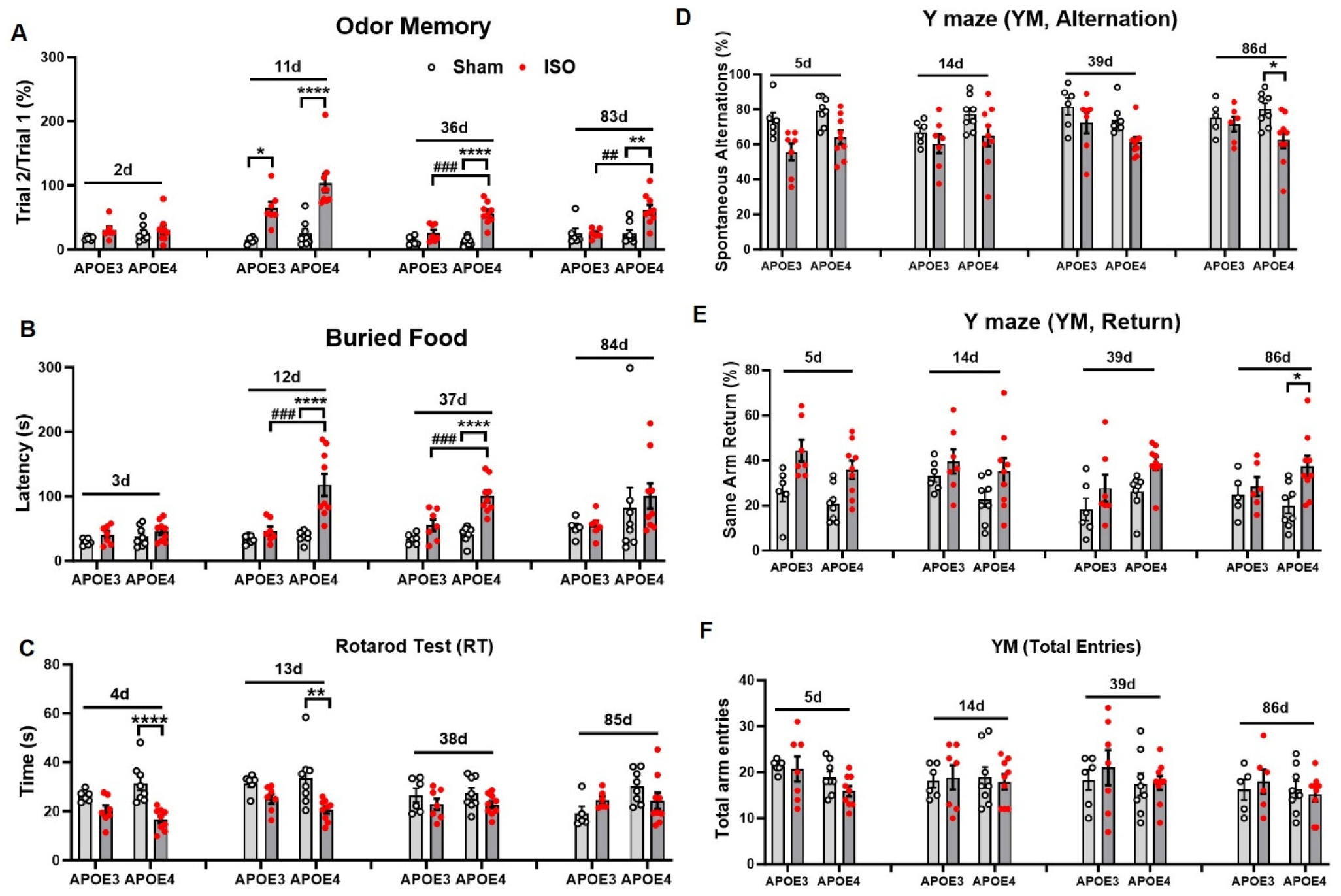
Isoflurane (ISO) exposure in E4 mice result in long-lasting neurological impairments. **A-B** Olfactory dysfunction in E4 mice was demonstrated using the odor memory (OM, A) and buried food (BF, B) tests. **C** E4 mice exhibited impaired neuromuscular and motor functions, as demonstrated by reduced performance in rotarod (RT) tests. **D-F** Spatial memory was assessed using the Y-maze test, which revealed a significant impairment in spontaneous alternations (D) and increased arm return (E) at day 86, while total arm entries remained unchanged (F). n=6 (E3/Sham), 7 (E3/ISO), 8 (E4/Sham), and 9 (E4/ISO). *p<0.05, **p<0.01, and ****p<0.0001 vs. E3/Sham. Data was analyzed with Two-way ANOVA with Tukey multiple comparisons test.

Next, we profiled OB transcriptomic changes 90 days after ISO/OP to assess long-term molecular effects in E4 mice. RNA-seq showed clear clustering by genotype and treatment (Fig. 7A; n=6 mice/group). Differential expression analysis with limma and FDR correction identified distinct differences between E4/ISO and E3/ISO mice (Fig. 7B). GO enrichment showed upregulation of pathways related to membrane potential regulation and calcium ion transport (Fig. 7C), and downregulation of metabolic, energetic, and protein-complex pathways (Fig. 7D). Representative genes from these pathways are shown in Fig. 7E-F. Lipid metabolism-related genes were also broadly reduced in E4/ISO mice, including chaperones (*Hspa8*, *Hsp90b1*, *Hsp90aa1*), stress-response genes (*Fos*, *Atf4*, *Xbp1*), and lipid/bioenergetic genes (*Cyp51*, *Hmgcs1*, *Msmol*; Fig. 7G-H). However, these ISO-associated changes were modest relative to baseline E3-E4 genotype differences, which showed stronger transcriptomic separation and lipid metabolism pathway differences (Supplementary Figure S13). In contrast, E4/ISO mice showed few OB DEGs compared with E4/Sham controls, whereas E3 mice exhibited more pronounced ISO-associated changes (Supplementary Figure S 14A-J). Lipid metabolism analysis identified 11 DEGs in the E3/ISO vs. E3/Sham comparison (Supplementary Figure S14K).

**Fig. 7.**
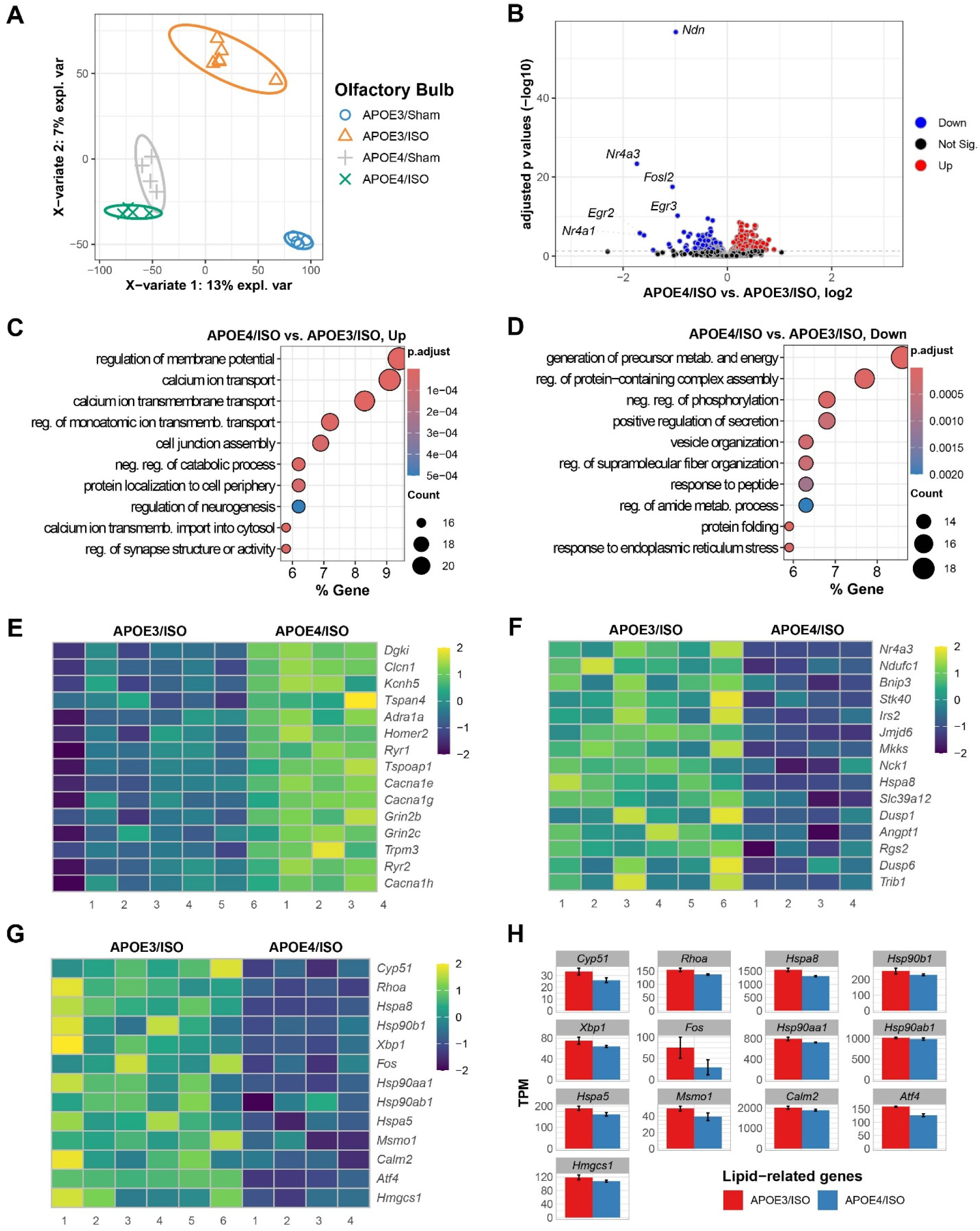
APOE4 drives long-term alterations to transcriptomic profile of OB region at 90d after isoflurane (ISO) exposure and laparotomy operation (OP). **A** PLS-DA plot for all normalized transcriptome genes is depicted for humanized E4 and E3 mice with or without ISO exposure. **B** Volcano plot displaying differentially expressed genes (DEGs) in the E4/ISO vs. E3/ISO comparison. **C-D** Pathway enrichment analysis of up (C) and downregulated (D) DEGs with Gene Ontology molecular processes. **E-F** Heatmaps displaying DEGs associated with the top three upregulated pathways (E) and top three downregulated pathways (F) identified between E4/ISO and E3/ISO groups. Z-score normalization was applied to gene expression values. Heatmaps illustrate sample-wise variation (columns) across genes (rows). Color intensity reflects relative transcript abundance across samples: warmer colors for higher, cooler for lower. **G-H** Heatmap and bar graph of genes involved in regulation of lipid metabolism. n=6 mice/group.

In the HI (Supplementary Figure S15), normalized counts also clustered by genotype and treatment, and E4/ISO differed from E3/ISO mice at FDR<0.05. Enrichment analysis showed upregulation of RNA splicing pathways and downregulation of organelle and ribosome assembly pathways in E4/ISO compared with E3/ISO mice. Baseline genotype differences in the HI were more robust than ISO effects, with E4/Sham vs. E3/Sham showing enrichment for circadian rhythm and chromatin remodeling among upregulated genes, and muscle contraction and NADH regeneration among downregulated genes (Supplementary Figure S16). No DEGs were detected in E4/ISO vs. E4/Sham, whereas E3/ISO vs. E3/Sham showed more DEGs, mainly involving downregulated adherent’s junction assembly and apical protein localization pathways, with *Actb* and *Ccn3* as representative genes. Lipid metabolism-focused analysis identified *Ubc* as the only related DEG in both E4/Sham vs. E3/Sham and E3/ISO vs. E3/Sham comparisons. Taken together, we found that ISO/OP induced long-lasting olfactory and late-emerging cognitive impairments specifically in APOE4 mice, alongside transcriptomic signatures in OB and HI that reveal disrupted metabolic, splicing, and ribosomal pathways. These effects highlight APOE4’s heightened vulnerability to anesthesia and surgery, even in asymptomatic young adults.

## 4 Discussion

By examining how short-term ISO/OP exposure interacts with the AD genetic risk factor APOE4, we identified APOE4-specific vulnerabilities in lipid metabolism, neural circuit function, and behavior. ISO/OP induced bioenergetic disruption in the OB and HI, glia-mediated lipid dysregulation, exacerbated neural circuit dysfunction, and persistent olfactory deficits that preceded later cognitive decline. Ninety days after ISO/OP exposure, transcriptomic analysis revealed downregulation of lipid metabolism and atherosclerosis-related pathways in E4 mice, suggesting accelerated molecular alterations associated with APOE4. Collectively, these findings indicate that ISO/OP exacerbates APOE4-associated neurobiological changes by disrupting glial lipid homeostasis, thereby impairing neural circuit integrity and contributing to progressive functional deficits.

Although surgical inflammation is a major contributor to postoperative cognitive dysfunction, anesthesia alone can also affect neural and behavioral outcomes. In our prior work with aged mice, ISO alone produced modest, transient changes, whereas combined anesthesia and surgery caused robust, sustained olfactory and cognitive impairments [7], supporting a synergistic effect. Thus, this study focused on the combined ISO/OP model, which better reflects the clinical perioperative setting. Although our transcriptomic data suggests immune pathway activation after ISO/OP, especially in aged animals [7, 41], the current study did not measure circulating or protein-level inflammatory biomarkers. Future studies incorporating peripheral and central immune readouts will help clarify the roles of inflammatory signaling and direct anesthetic effects. Lipids as essential nutrients for all living organisms play a critical physiological role in the brain, the second most lipid-rich organ after the adipose tissue where they account for 10-12% of the brain’s fresh weight and over 50% of its dry weight [42]. In AD, aggregation of lipid droplets, dysfunctional lipid metabolism and increased levels of lipid oxidation have been recognized as an important hallmark of neuropathology alongside amyloid plaques and neurofibrillary tangles [43, 44]. In APOE4 carriers, lipid signatures of both neurons and glial cells include increased levels of Cer and FA [45–47], leading to increased risk for AD. Our findings built on these previous studies have provided evidence of additive effects of ISO/OP on prodromal neuronal and glial pathology, contributing to the development of PND. Both OB and HI-derived glial lipidomic profiling revealed that, even in young asymptomatic mice, short-term exposure to ISO/OP unmasked E4-specific functional deficits and lipid perturbations. Despite multiple prior studies reporting the effects of isoflurane and other anesthetics on lipid composition and peroxidation [48–50], our study presented the first demonstration of E4 and isoflurane interaction to impact lipid regulation in the brain.

Specifically, the depleted levels of TG in the OB of E4/ISO mice compared to E3 counterparts signifies either increased lipolysis or impaired synthesis. Since TG is the storage form of FA, its depletion may indicate reduced capacity to buffer excess FA, which corresponds to the increased levels of MDA observed in the same assay. Another notable result derived from lipid composition was the changes in E4/ISO mice compared to E4/Sham group, which featured high levels of FA and MDA in the OB region, alongside lower levels of CL in the hippocampus. Since CL is a mitochondrial signature lipid essential for maintaining cristae structure, stabilizing respiratory chain complexes, and supporting oxidative phosphorylation, its depletion suggests impaired mitochondrial bioenergetics. Consistently, reductions in CL have been previously associated with mitochondrial dysfunction in AD and related disorders [51]. These observations are further supported by lipidomic results derived from microglia and astrocytes, which indicated remodeling of membrane composition, altered mitochondrial integrity and impaired lipid droplet buffering after isoflurane.

Our lipidomic and physiological findings support the notion that OD may signal anesthesia- and E4-related vulnerability, appearing before overt cognitive decline. The OB, being highly metabolically active and among the earliest brain regions to exhibit E4-driven pathology [52], is particularly susceptible to additional metabolic stressors such as general anesthesia and surgery. We observed lipid dysregulation in the OB, including TG depletion, phospholipid remodeling, and accumulation of bioactive sphingolipids, suggesting impaired lipid storage and trafficking capacity in glial cells, which normally buffer neurons against energetic fluctuations. Concomitant CL loss and altered phosphatidylethanolamine composition point to mitochondrial stress and compromised oxidative phosphorylation, consistent with a shift toward bioenergetic insufficiency. Isoflurane’s effects on neuronal excitability and synaptic balance were strongly modulated by APOE genotype, largely due to lipidomic dysregulation in astrocytes and microglia that are critical regulators of brain homeostasis [53, 54]. Astrocytes, the brain’s primary source of APOE, are responsible for cholesterol and phospholipid transport to neurons [53, 55]. In APOE4 carriers, deficits in astrocytic lipid handling impair membrane maintenance, and exposure to isoflurane likely worsens these disruptions, altering neuronal and glial membrane composition and affecting synaptic function. Microglia, key players in synaptic pruning and inflammation, are also sensitive to lipid imbalance. Isoflurane-induced changes in lipids like ceramides and sphingolipids can activate microglia, particularly in E4 mice where lipid turnover is already impaired [56]. This may underline the observed mitral cells hypoexcitability and disrupt gamma oscillations in E4 mice. Conversely, E3 astrocytes and microglia may more effectively restore lipid homeostasis following isoflurane exposure, leading to transient hyperexcitability without long-term circuit dysfunction. Collectively, these findings suggest that OD reflects not only local circuit vulnerability but also serve as an early, functionally accessible readout of APOE4- and anesthesia-associated disruptions in lipid metabolism and neuronal-glial energetics.

A notable discrepancy in our findings was the robust lipidomic remodeling at 7 days post-ISO/OP compared with the relatively few DEGs detected by bulk RNAseq. This divergence likely reflects the fact that lipid metabolism is extensively regulated at post-transcriptional and enzymatic levels, enabling rapid remodeling without corresponding shifts in gene expression. In addition, bulk RNA-seq averages signals across heterogeneous cell types, potentially masking subtle but functionally important changes confined to astrocytes or microglia. By contrast, lipidomic analysis provides a more sensitive snapshot of metabolic state, capturing perturbations in energy storage and membrane composition that may precede broader transcriptional reprogramming.

Building on Foley et al.’s report of distinct E4-driven cortical gene signatures in young mice [57], our RNA-seq analysis revealed divergent transcriptional responses to ISO/OP in E3 and E4 mice at 90 days. E3 mice showed broad changes in pathways related to metabolism, protein complexes, and calcium signaling, suggesting active molecular remodeling after anesthesia and surgery. In contrast, E4 mice showed few DEGs after ISO/OP despite clear functional impairments. This muted response may reflect a pre-stressed or primed state [58–60], in which baseline alterations in lipid metabolism, stress response, and chaperone activity limit further transcriptional adaptation. Alternatively, because E4 mice can develop OD by 6-8 months of age [61], spontaneous pathology in age-matched E4/Sham mice may have reduced detectable differences from E4/ISO mice. Together, these findings suggest that E4 may confer impaired transcriptomic plasticity after perioperative stress, contributing to persistent olfactory deficits and delayed cognitive vulnerability.

A previous study showed that anesthesia/surgery induces olfactory and cognitive impairment and proposed an IL-6–mediated pathway linking olfactory dysfunction to hippocampal changes [4]. In contrast, our study examines this relationship in APOE4 mice and identifies distinct vulnerabilities at the level of glial lipid metabolism and olfactory circuit function. Our findings further suggest that olfactory and circuit-level alterations emerge early following perioperative exposure, preceding overt cognitive deficits.

As with other humanized APOE knock-in studies, species-specific differences and the use of 100% oxygen during anesthesia should be considered when extrapolating these findings to clinical populations. Another limitation is the use of male-only cohorts. Although sex differences exist, E4-associated phenotypes are observed in both sexes, with sex often modulating the magnitude or trajectory of these effects rather than their presence [62–65]. Male mice were selected as an initial cohort to determine whether E4 confers susceptibility to perioperative glial lipid dysregulation and neurobehavioral deficits relative to E3. Direct comparison of male and female responses to perioperative ISO exposure remains an important next step, particularly given evidence that E4-related metabolic and cognitive phenotypes may be more pronounced in females.

From a translational standpoint, these findings support OD assessment, network dysfunction, and lipidomic profiling as potential early markers of perioperative vulnerability, particularly in E4 carriers. Routine olfactory testing and circulating lipidomic signatures, including extracellular vesicle-derived lipids, may help identify patients at higher risk for PND or accelerated neurodegeneration and inform clinical trial stratification or personalized perioperative monitoring. Because this study focused on isoflurane, future work should determine whether these lipid and circuit disruptions extend to other anesthetic agents. Interventions that stabilize lipid homeostasis or support mitochondrial function may also offer strategies to mitigate perioperative risk in genetically vulnerable patients.

In summary, our multidisciplinary approach demonstrates that E4 mice are particularly vulnerable to perioperative stress, with ISO/OP accelerating glial lipid dysregulation, mitochondrial dysfunction, circuit instability, and behavioral deficits. These changes emerged at metabolic and circuit levels before broad transcriptional remodeling, supporting lipidomic changes and olfactory dysfunction as early indicators of vulnerability. By linking glial lipid metabolism to circuit and behavioral outcomes, our study highlights APOE genotype as an important factor in perioperative neurological risk and provides a framework for future studies of anesthetic choice, patient stratification, and targeted intervention.

## Abbreviations

AD: Alzheimer’s disease
APOE: Apolipoprotein E
aCSF: artificial cerebrospinal fluid
BSA: Bovine serum albumin
BF: Buried food
CL: Cardiolipins
DAPI: 4,6-diamidino-2-phenylindole
DEGs: Differentially expressed genes
ETCs: External tufted cells
E3: Apolipoprotein E3
E4: Apolipoprotein E4
FDR: False discovery rates
FA: Fatty acid
GA: General anesthesia
GO: Gene ontology
GS: Grip strength
HBSS: Hank’s balanced salt solution
HexCer: Hexosylceramides
HI: Hippocampus
ISF: Instantaneous spike frequency
ISO: Isoflurane
LC-MS/MS: Liquid chromatography coupled with tandem mass spectrometry
LFP: Local field potential
LPC: Lysophosphatidylcholine
LPE: Lysophosphatidylethanolamine
MACS: Magnetic-activated cell sorting
MDA: Malondialdehyde
MCL: Mitral cell layer
NB: Nest building
NMDG: N-Methyl-D-glucamine
NOR: Novel object recognition
OB: Olfactory bulb
OM: Odor memory
OD: Olfactory deficits
OF: Open field
OP: Operation
PCA: Principal component analysis
PB: Phosphate buffer
PC: Phosphatidylcholines
PE: Phosphatidylethanolamine
PND: Perioperative neurocognitive dysfunction
PNs: Pyramid neurons
PSD: Power spectrum density
RT: Rotarod test
RNAseq: RNA sequencing
SM: Sphingomyelin
sEPSCs: Spontaneous excitatory postsynaptic currents
sIPSCs: Spontaneous inhibitory postsynaptic currents
TPM: Transcripts per million
TG: Triglycerides
YM: Y maze

## Supplementary Information

The online version contains supplementary material and figures available at xxxx.

## Acknowledgements

The confocal imaging data used in this publication was produced in collaboration with the Biomedical Microscopy Core at the University of Georgia. We thank Miss. Anna Wu Chen at UCSD for her substantial editing of the manuscript.

## Authors’ contributions

JW and SL conceived the project and designed the experiments. YL, YJ, CU, STI, MH, DZ, ZW, and HL performed experiments and sample processing. YL, YJ, CU, STI, MH, DZ, JWJ and HL performed data curation, formal analysis, investigation, and visualization. YL, YJ, SL, and JW wrote the manuscript. All authors approved the final version of the manuscript.

## Data availability

All data supporting the conclusions of this study are provided in the main text and/or in the Supplementary Materials, which include 16 supplemental figures with their corresponding legends.

## Declarations

### Ethics approval and consent to participate

All experimental procedures were approved by the Institutional Animal Care and Use Committees (IACUC) of the University of Maryland School of Medicine and the University of Georgia.

### Consent for publication

Not applicable.

