## Supplemental Materials for "Isoflurane and surgery aggravate APOE4-dependent lipid dysregulation and neural dysfunction, leading to neurological impairment in male mice"

**Supplemental Figures (16)**

**Supplemental Figure Legends**

**Supplemental Materials**

**Materials and Methods**

**Mouse general anesthesia and operation**

Homozygous humanized APOE4 KI [B6(SJL)-Apoetm1.1($^{APOE*4}$)Adiuj/J, stock no. 027894] and APOE3 KI [B6.Cg-Apoeem2$^{(APOE*)}$Adiuj/J, stock no. 029018] mice (Mus musculus; male; 10-14 weeks old; 20-25 g) were obtained from The Jackson Laboratory. These knock-in mice carry targeted replacement of the endogenous mouse *Apoe* gene with human *APOE* sequences. Animals were maintained under specific pathogen-free conditions and housed under a 12-hour light/dark cycle with ad libitum access to food and water. GA was induced and maintained with 2% ISO in 100% oxygen via a precision vaporizer. Following induction, animals were transferred to a surgical station and maintained under continuous ISO administration via a nose cone. During the surgical phase, anesthetic depth was assessed by monitoring respiratory rate and pattern, as well as palpebral and pedal reflexes. Core body temperature was maintained using a temperature-regulated heating pad. A midline laparotomy was performed through a 1.5 cm incision extending from the xiphoid process to 0.5 cm above the pubic symphysis, sequentially incising the skin, abdominal musculature, and peritoneum. At the conclusion of the procedure, the incision site was infiltrated with 0.25% bupivacaine in sterile saline and closed in layers using 5-0 monofilament nylon sutures. The surgical procedure lasted approximately 10 minutes. Following surgery, animals were returned to the anesthesia chamber and maintained under 2% ISO to complete a total exposure time of 2 h. During this period, respiratory rate and pattern were continuously monitored to ensure stable anesthetic depth, and no signs of respiratory distress or instability were observed. All animals were maintained under identical anesthetic conditions.

Postoperatively, mice were placed on a temperature-controlled pad for 30-60 minutes to facilitate thermal recovery. Animals were closely monitored for 4 h following general anesthesia and surgery, with continued daily assessments thereafter.

**Adult cell isolation**

Dissected OB and hippocampus (HI) were collected in ice-cold Hank's Balanced Salt Solution (HBSS) and enzymatically dissociated into single-cell suspensions using the Adult Brain Dissociation Kit (Cat# 130-107-677, Miltenyi Biotec), following the manufacturer's protocol. Homogenates were passed through a 70-μm cell strainer to remove debris and washed with HBSS supplemented with 0.5% bovine serum albumin (BSA). Myelin and cellular debris were eliminated by density gradient centrifugation with the Debris Removal Solution. The resulting suspension was resuspended in MACS buffer (PBS, 0.5% BSA, 2 mM EDTA) and incubated with

FcR Blocking Reagent to minimize nonspecific binding. For microglial isolation, cells were labeled with CD11b Microbeads (Cat# 130-093-634, Miltenyi Biotec), whereas astrocytes were enriched in parallel using ACSA-2 Microbeads (Cat# 130-097-678, Miltenyi Biotec). Magnetically labeled cells were applied to LS separation columns placed in a MACS Separator (Miltenyi Biotec). After successive washes, positively selected CD11b+ microglia and ACSA-2+ astrocytes were eluted.

**Lipidomics**

Cell isolation and subsequent lipidomic assay was performed on pooled samples obtained from 3 male mice (10-14 weeks old) of the same genotype (APOE3/APOE4) and treatment group (Sham/ISO) at 7 d after cessation. Briefly, total lipids were extracted using the MTBE protocol and analyzed by liquid chromatography coupled with tandem mass spectrometry (LC-MS/MS). The LC-MS/MS analyses were conducted on an Ultimate 3000 Ultra High-Performance Liquid Chromatograph coupled to a Thermo TSQ Altis Triple Quadrupole Mass Spectrometer (Thermo Scientific, San Jose, CA). Chromatographic separation was achieved using an ACQUITY Amide BEH column (1.7 μm, 2.1 × 100 mm; Waters, Milford, MA) maintained at 45 °C. The mobile phases consisted of solvent A (ACN/$H_2O$, 95:5, v/v) and solvent B (ACN/$H_2O$, 50:50, v/v), each containing 10 mM ammonium acetate. The gradient program was run at 0.6 mL/min with the following profile: 0.1-20% B over 2 min, 20-80% B over 3 min, 80% to 0.1% B in 0.1 min, followed by equilibration at 0.1% B for 2.9 min. The total run time was 8.0 min, with a 2 μL injection volume. The autosampler was maintained at 7 °C. Electrospray ionization was performed in both positive and negative ion modes, and mass spectrometric detection employed SRM mode. ESI source settings were spraying voltage of +3500 V (positive mode) and −2500 V (negative mode), sheath gas (Arb) = 60, auxiliary gas (Arb) = 15, sweep gas (Arb) = 1, and ion transfer tube temperature of 380 °C. Nitrogen was used as the nebulizer gas and argon as the collision gas (1.5 mTorr). The vaporizer temperature was 350 °C. Collision energies and RF lens voltages were optimized for each lipid class reference standard.

***In vivo* electrophysiological recording and data analysis**

Animals were anesthetized by 2% ISO. To prepare for recordings from the mitral cell layer (MCL) on the medial side of each OB of awake mice, the craniotomy was made on the dorsal skull with the following coordinates: 5.50 mm anterior from Bregma and 0.30 mm lateral from the midline. A metal head-plate with a 4.2-mm round opening (Models 1 and 10; Neurotar, Helsinki, Finland) was attached to the skull with Metabond. One week after surgery, mice were trained for habituation to head fixation in head-fixed treadmill apparatus (Neurotar, Helsinki, Finland) for 15 min/day in 5 consecutive days. On day 13 after headplate implantation, animals received a single 2 h ISO exposure as the experimental treatment. ISO was delivered at 5% for induction and 2% for maintenance with a room air flow rate of 75 ml/min. The depth of anesthesia was confirmed by the loss of hind paw withdrawal reflex tested every 15 minutes. This paradigm isolates the effects of ISO anesthesia following prior cranial surgery required for recording. Groups compared include E3/Sham, E4/Sham, E3/ISO, E4/ISO.

To verify the position of the recording sites, probes were coated with two different dyes: DiI-DiIC18(3) (red) for the left OB and DiO-DiOC18(3) (green) (Invitrogen) for the right OB. Electrophysiological signals were detected by the neural probe and passed through a digital head stage to a 32-channel amplifier (Plexon DigiAmp, Plexon, TX, USA), where they were sampled at 1 kHz and bandpass filtered at 300–7,500 Hz for spikes and at 0.1-200 Hz for local field potential (LFP) before being digitally sampled at 40 kHz by a Plexon Omniplex recording system (Plexon, Dallas, TX).

**Patch clamp recording in brain slices and data analysis**

Animals were deeply anesthetized with ISO using an open-drop method prior to decapitation. Horizontal OB and coronal brain sections were cut using a VT1200S vibratome (Leica Microsystems, Germany) in ice-cold, oxygenated (95% $O_2$/5% $CO_2$) N-Methyl-D-glucamine (NMDG)-based artificial cerebrospinal fluid (aCSF) containing (in mM): 90 NMDG, 2.5 KCl, 1.2 NaH2PO4, 30 NaHCO3, 25 glucose, 20 NaHEPES, 5 Sodium ascorbate, 2 Thiourea, 3 Sodium pyruvate, 10 MgSO4, 0.5 CaCl2 (pH 7.40, 300 mOsm). After 30 min of incubation in NMDG aCSF at 30°C, slices were then transferred to HEPES-based aCSF at RT (room temperature) until they were used for recordings. HEPES-based aCSF was continuously bubbled with 95% O2-5% CO2 and had the composition (in mM): 92 NaCl, 2.5 KCl, 1.2 NaH2PO4, 30 NaHCO3, 25 glucose, 20 NaHEPES, 2 Sodium ascorbate, 2 Thiourea, 3 Sodium pyruvate, 2 MgSO4, 2 CaCl2 (pH 7.40, 300 mOsm). During the experiments, slices were perfused at 3 ml/min with recording aCSF, which was equilibrated with 95% O2/5% CO2 and warmed to 30°C and had the following composition (in mM): 125 NaCl, 2.5 KCl, 1.25 NaH2PO4, 25 NaHCO3, 10 glucose, 2 MgSO4, 2 CaCl2 (pH 7.40, 300 mOsm).

Whole-cell patch-clamp recordings were performed from neurons in OB or HI slices visualized with an Eclipse FN1 fixed-stage upright microscope (Nikon, Japan) equipped with near-infrared differential interference contrast optics. External tufted cells in the OB slices or pyramidal neurons in the CA1 region of HI slices were preselected for recording based on their soma location, sizes and shapes as well as the projection patterns of their primary dendrites. Cell identities were further verified by electrophysiological signatures and post hoc morphological reconstruction with histocytochemistry. Biocytin (0.2%) was included in the patch pipette internal solution to fill the recorded neurons thus enable post hoc visualization of their somatodendritic architectures. Electrophysiological signals were acquired using the MultiClamp 700B amplifier (Molecular Devices) in current- or voltage-clamp and pClamp Data Acquisition Software (Clampex v11). Data were sampled 2 kHz or 10 kHz and filtered at 5 kHz or 50 kHz in voltage or current clamp, respectively.

**Histochemistry for neural probe position verification and neuronal reconstruction**

For histological verification of the 32-channel neural probe positioning in the OB, the mouse brain was dissected immediately after *in vivo* recording and fixed in 4% paraformaldehyde overnight at 4°C. The OB tissue was cut into 150 µm thick coronal sections using a Compresstome® Vibrating Microtomes (VF-510-0Z, Precisionary Instruments, Ashland, MA). Then brain sections were washed in 0.1 M phosphate buffer (PB) for 3 times (5 minutes/each) at room temperature before being incubated in 0.25% Triton X-100 in 0.1 M PB for 1 hour at room temperature for permeabilization. After 3 times (5 minutes/each) wash with 0.1 M PB, brain sections were stained with 4,6-diamidino-2-phenylindole (DAPI) (1:200 dilution) for 10 minutes to label cell nuclei at room temperature and protected from light. DAPI binding was terminated by another 3 times (5 minutes/each) wash with 0.1 M PB before brain sections were mounted onto microscope glass slides using anti-fade mounting medium.

For morphological reconstruction of the patch clamp recorded neurons in the OB or HI, brain slices with biocytin (0.2%, w/v)-filled cells were immediately kept in 4% paraformaldehyde at 4°C overnight. After three (5 min each) washes with 0.05 m phosphate-buffered saline (PBS), slices were incubated in a blocker solution on a shaker for 1 h. Blocker solution was made by 0.05 m PBS with the addition of bovine serum albumin and Triton X-100 with final concentrations of 1% (w/v) and 0.5% (v/v), respectively. The slices were then transferred to and kept in this blocker solution containing streptavidin-CY3 (1 µg/mL) covered with aluminum foil to prevent light exposure at room temperature on a shaker for 7 h. Following three (5 min each) rinses with 0.05 m PBS to terminate streptavidin-CY3 staining, slices were treated with 0.05 m PBS containing DAPI (5 µg/mL) at room temperature in the dark for 10 min. DAPI staining was terminated by three (5 min each) washes with PBS before slices was wet mounted and cover-slipped with fluorescence mounting media. Confocal images were acquired using a Zeiss LSM 900 microscope equipped with a 2.5× or 40× oil-immersion objective. Z-stacks were collected with a step size of 1 µm and an XY resolution of 0.21 µm/pixel.

**Neurological assessment**

Odor memory (OM) test: To evaluate olfactory learning and memory, mice were singly housed overnight and subjected to a previously described OM protocol [1, 2]. A sterile, 6-inch cotton-tipped wooden applicator was introduced through a cage-lid port, allowing the animal 30 minutes of familiarization. Each mouse was provided with a new applicator. For odor presentation, cinnamon powder (McCormick; 100 ng/ml in water) was freshly prepared and stored in airtight vials. During the first trial (T1), the applicator tip was briefly immersed in the odor solution (2 s) and inserted ~2.5 cm into the cage. Sniffing behavior, defined as nose orientation toward the applicator within 2 cm, was recorded over 5 minutes. Following a 60-minute interval, a second trial (T2) was conducted identically. Memory retention was quantified as the [T2/T1] × 100 sniffing ratio, with reduced exploration in T2 reflecting recognition of the familiar odor.

Buried food (BF) test: To assess olfactory function, BF was performed to measure the animal's ability to detect and retrieve a familiar food reward hidden under bedding as described in previous papers [1, 3]. Mice were individually housed with free access to water but food-deprived for 24 h to increase motivation. The evening prior to testing, each subject received a mini cookie in its home cage for odor familiarization, and consumption was confirmed the next morning. On the test day, mice were acclimated for 10 min in a clean cage (46 × 23.5 × 20 cm) containing 3 cm of fresh bedding, then returned to their home cage. A cookie (~1 g) was buried 2-3 cm beneath the bedding in a randomly chosen corner, and the mouse was reintroduced. The latency to locate and begin eating the buried cookie was recorded, with trials ending after 15 min if unsuccessful.

Nest building (NB) test: The NB is a widely used non-invasive behavioral assay, was employed to assess rodent well-being, motivation, and cognitive function [4]. Mice were singly housed in standard cages containing wood chip bedding. Each cage was supplied with a pre-weighed pressed cotton square (nestlet) as the sole nesting material. Animals were left undisturbed overnight (~12 h), after which nest quality was scored the next morning on a standardized 5-point scale, ranging from 0 (nestlet untouched) to 5 (well-formed dome). Remaining nesting material was weighed to quantify usage, and nests were photographed for documentation. Poor performance in this task is indicative of deficits in executive function, motor planning, or motivation.

Grip strength (GS) test: This test in mice is designed to measure neuromuscular function, particularly forelimb and/or hindlimb strength using a digital grip strength meter (Ugo Basile) based on our established protocols[3]. Briefly, the forelimb GS of both paws was assessed by placing the mouse on a mesh wire grid connected to the device's force transducer. Once the mouse secured a firm grip, it was gently held by the tail and slowly pulled away from the grid. The maximum force exerted on the mesh wire grid was recorded, with each mouse undergoing an average of 10 daily trials. Final GS values were normalized to body weight for comparison between groups.

Rotarod (RT) test: Locomotor performance was assessed using the Rotarod (Harvard Apparatus) following established protocols[5]. The device was programmed to accelerate from 4 to 40 rpm over 90 s, with each trial capped at 300 s. Mice were positioned on the rotating rod prior to start, and the latency to fall was recorded. For each animal, the mean latency across five trials was calculated and used for group comparisons.

Y maze (YM): The YM test was used to assess hippocampus-dependent spatial working memory in mice, following established methods [3]. The apparatus (Stoelting Co.) consisted of three identical arms (A, B, C). Each trial began with the mouse placed at a randomly chosen start arm, after which it was allowed to explore freely for 5 min. Arm entries were recorded, and a spontaneous alternation was defined as consecutive entries into three different arms. Alternation percentage was calculated as: (total alternations ÷ [total arm entries – 2]) × 100. Mice achieving scores above 50%, the chance level, were considered to have intact spatial working memory.

Open field (OF) test: Spontaneous locomotor activity was assessed in the OF test under dim red illumination[3, 6]. Mice were placed in a corner of the 22.5 × 22.5 cm chamber, facing the wall, and allowed to explore for 5 min. Movements were tracked using the Any-maze software, which quantified parameters such as total distance traveled, mean velocity, immobility duration, and percentage of time spent in the central zone.

Novel object recognition (NOR) test: Non-spatial recognition memory was assessed using the NOR test, which exploits the rodent's innate preference for novelty [3]. On day 1, mice were habituated to the open-field arena for 5 min. On day 2, they were exposed to two identical objects, and exploration time for each was recorded with ANY-maze software (Stoelting) until a total of 30 s was reached (maximum session time: 40 min). On day 3, one familiar object was replaced with a novel one, and testing proceeded under the same conditions. Recognition memory was considered intact when mice spent >15 s exploring the novel object.

**RNAseq analysis**

We used STAR aligner version 2.7.5 to map RNAseq reads to the mouse reference genome (GRCm39) with GENCODE gene annotation (version M33) [7]. We quantified gene and isoform expression at the transcripts per million (TPM) level using RSEM 1.3.3 [8]. We used DESeq2 version 1.42.1 for the differential expression analysis, with false discovery rates (FDR) derived from the Benjamini-Hochberg method [9]. Genes with an FDR of less than 0.05 were considered to be differentially expressed genes (DEGs) and used for downstream pathway enrichment analysis. Gene ontology (GO) analysis was performed with cluster Profiler 4.10.1 using DESeq2 output [10].

**Additional statistical and experimental design details**

Distinct animal cohorts were used for different experimental modalities unless otherwise specified. Because tissue collection required euthanasia, data obtained from separate tissue, cellular, electrophysiological, and molecular experiments represent independent biological samples. RNA sequencing was performed using tissue collected from mice after completion of behavioral testing, whereas electrophysiological recordings and most other assays were conducted in independent cohorts.

Sample sizes were based on prior studies and pilot experiments using comparable experimental designs and endpoints. Group sizes were selected to detect biologically relevant differences and were generally sufficient to detect medium-to-large effects at $\alpha = 0.05$ according to Cohen's effect-size conventions. No formal prospective power calculation was performed because the study incorporated multiple exploratory and mechanistically distinct experimental modalities.

Animals were included according to the prespecified genotype, age, sex, treatment, and experimental-completion criteria described in the relevant Methods subsections. All animals that completed the experimental protocol were included in the analysis. No animals or data points were excluded based on the observed outcome. Humane endpoints were established a priori in accordance with institutional IACUC guidelines; no animals met these criteria during the study.

Outcome measures included performance across multiple behavioral assessments, electrophysiological properties such as membrane resistance, resting membrane potential, synaptic transmission, and current injection–evoked firing, and molecular, transcriptomic, and lipidomic profiles of brain tissues and isolated cell populations. Individual outcome measures are described in the corresponding Methods subsections and summarized in the experimental workflow schematic (Supplementary Fig. 1). Given the exploratory, multimodal design of the study, no single primary outcome measure was predefined.

For each experiment, the specific statistical model, factors, post hoc comparisons, multiple-testing corrections, and sample sizes are reported in the corresponding figure legends or assay-specific Methods subsections. Statistical significance was defined as $p < 0.05$ unless otherwise specified. For high-dimensional analyses, adjusted p values or false-discovery-rate thresholds were applied as described for each dataset.

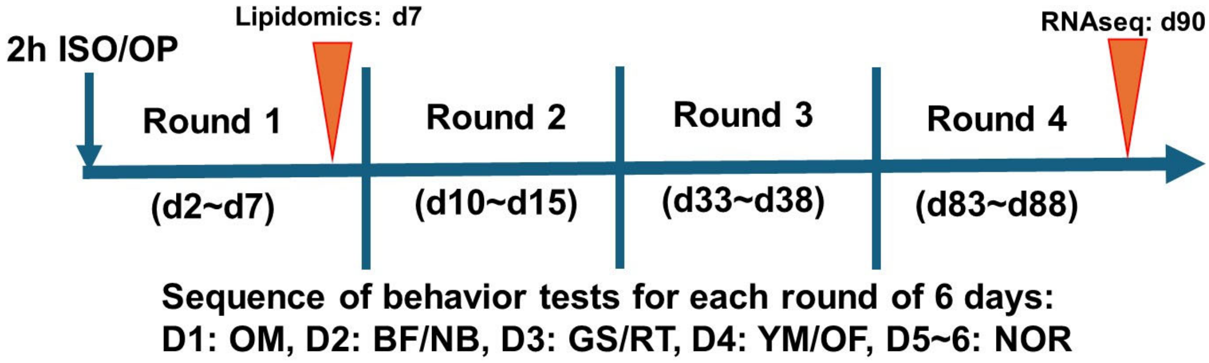


**Fig. S1. Schematic illustration of the experimental workflow for a battery of neurological behavioral tests and timing of tissue collection.** In addition to the cohort of mice used for evaluation of neurological functions, a separate age-matched cohort underwent the same treatment and was used for lipidomics. At 90d after cessation, the hippocampus and olfactory bulb regions were extracted from the behavioral study cohort and used for RNA sequencing (RNAseq). OM: odor memory, BF: buried food, NB: nest building, GS: grip strength, RT: rotarod, YM: Y maze, OF: open field, NOR: novel object recognition.

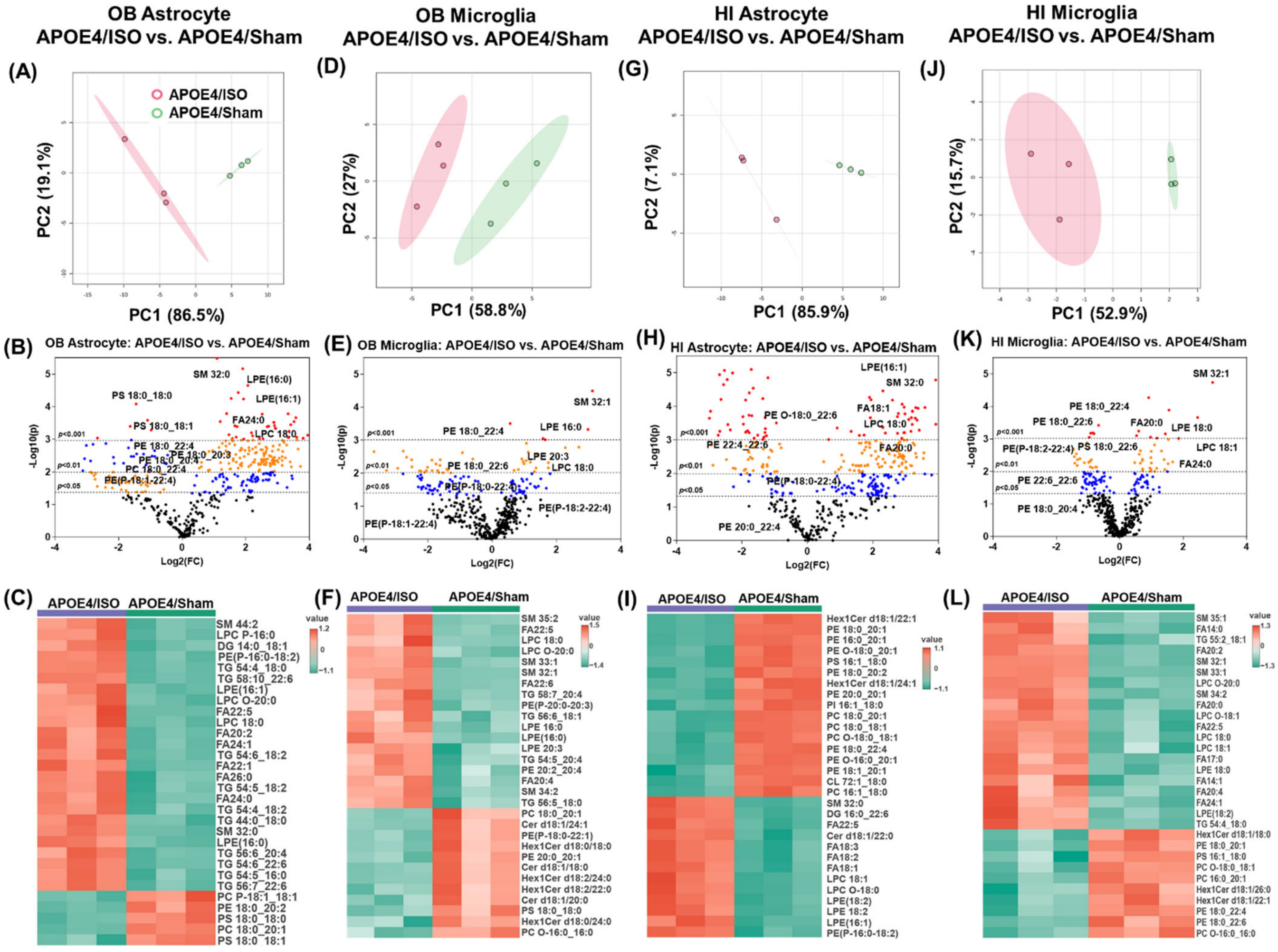


**Fig. S2. Isoflurane (ISO) exposure induces lipid dysregulation in the olfactory bulb (OB) and hippocampus (HI) astrocytes and microglia of APOE4 mice. (A–C)** OB astrocytes, **(D–F)** OB microglia, **(G–I)** hippocampal astrocytes, and **(J–L)** hippocampal microglia from APOE4/ISO vs. APOE4/Sham mice at 7 days post-exposure. In each set, PCA illustrates group separation, with volcano plots and heatmaps depicting differentially abundant lipids. n=3 samples/group. Each sample was pooled from three male mice, with a total of nine mice per group.

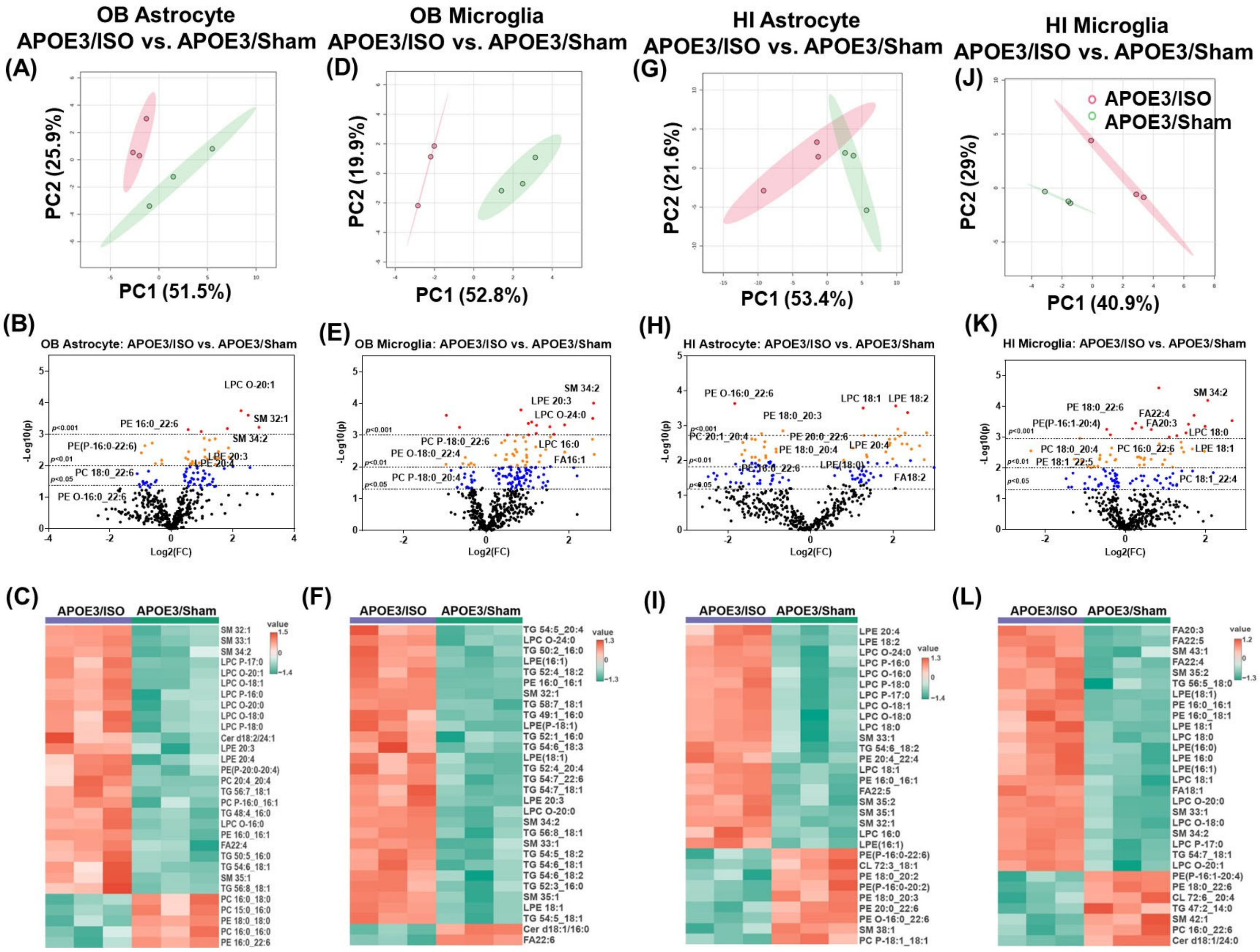

**Fig. S3. Isoflurane (ISO) exposure induces olfactory bulb (OB) and hippocampus (HI) glial lipid dysregulation, even in APOE3 mice. (A–C)** OB astrocytes, **(D–F)** OB microglia, **(G–I)** hippocampal astrocytes, and **(J–L)** hippocampal microglia from APOE4/ISO vs. APOE4/Sham mice at 7 days post-exposure. In each set, PCA illustrates group separation, with volcano plots and heatmaps depicting differentially abundant lipids. n=3 samples/group. Each sample was pooled from three male mice, with a total of nine mice per group.

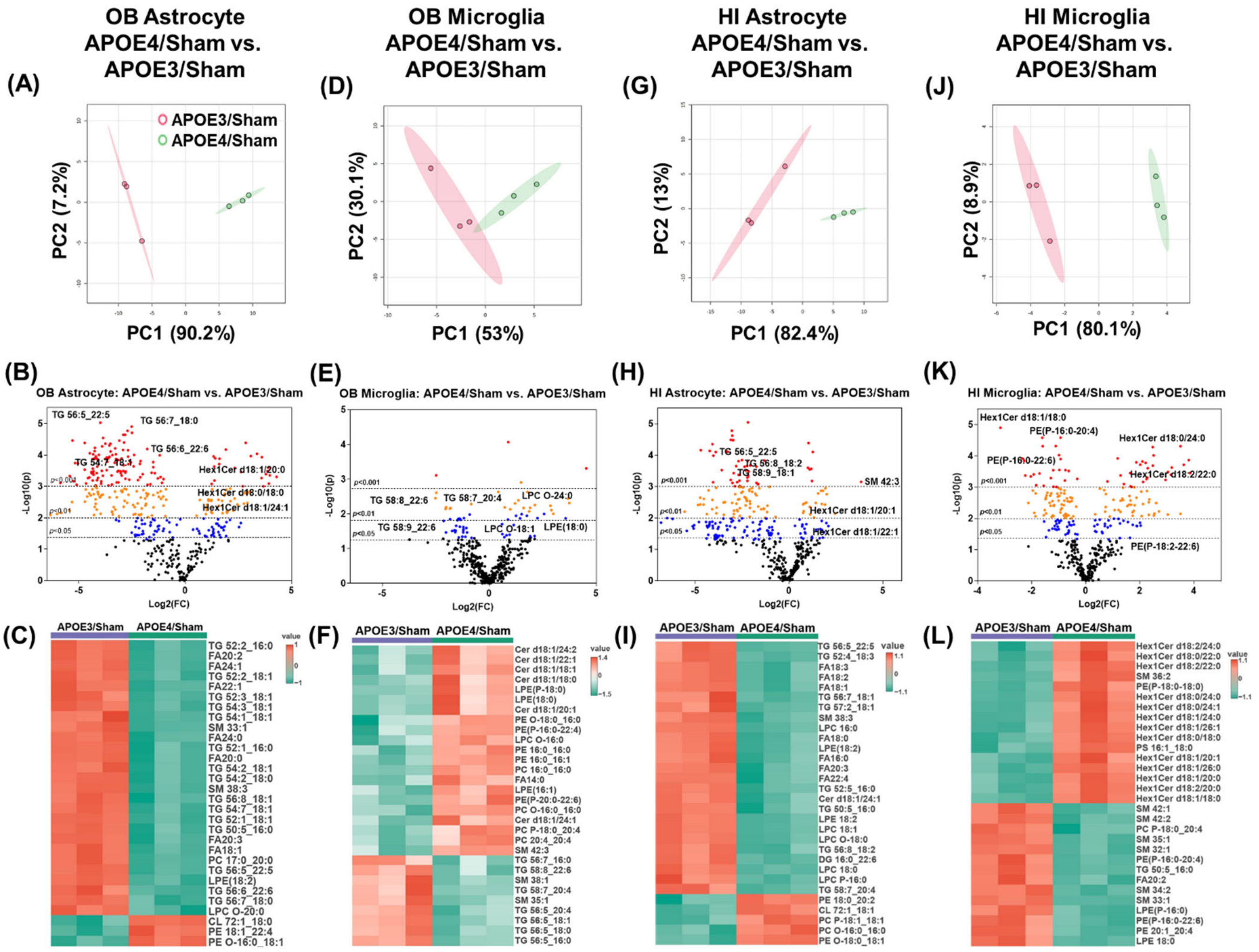


**Fig. S4. Humanized APOE4 mice display distinct baseline glial lipid profiles. (A–C)** OB astrocytes, **(D–F)** OB microglia, **(G–I)** hippocampal astrocytes, and **(J–L)** hippocampal microglia from APOE4/Sham vs. APOE3/Sham mice. In each set, PCA illustrates group separation, with volcano plots and heatmaps depicting differentially abundant lipids. n=3 samples/group. Each sample was pooled from three male mice, with a total of nine mice per group.

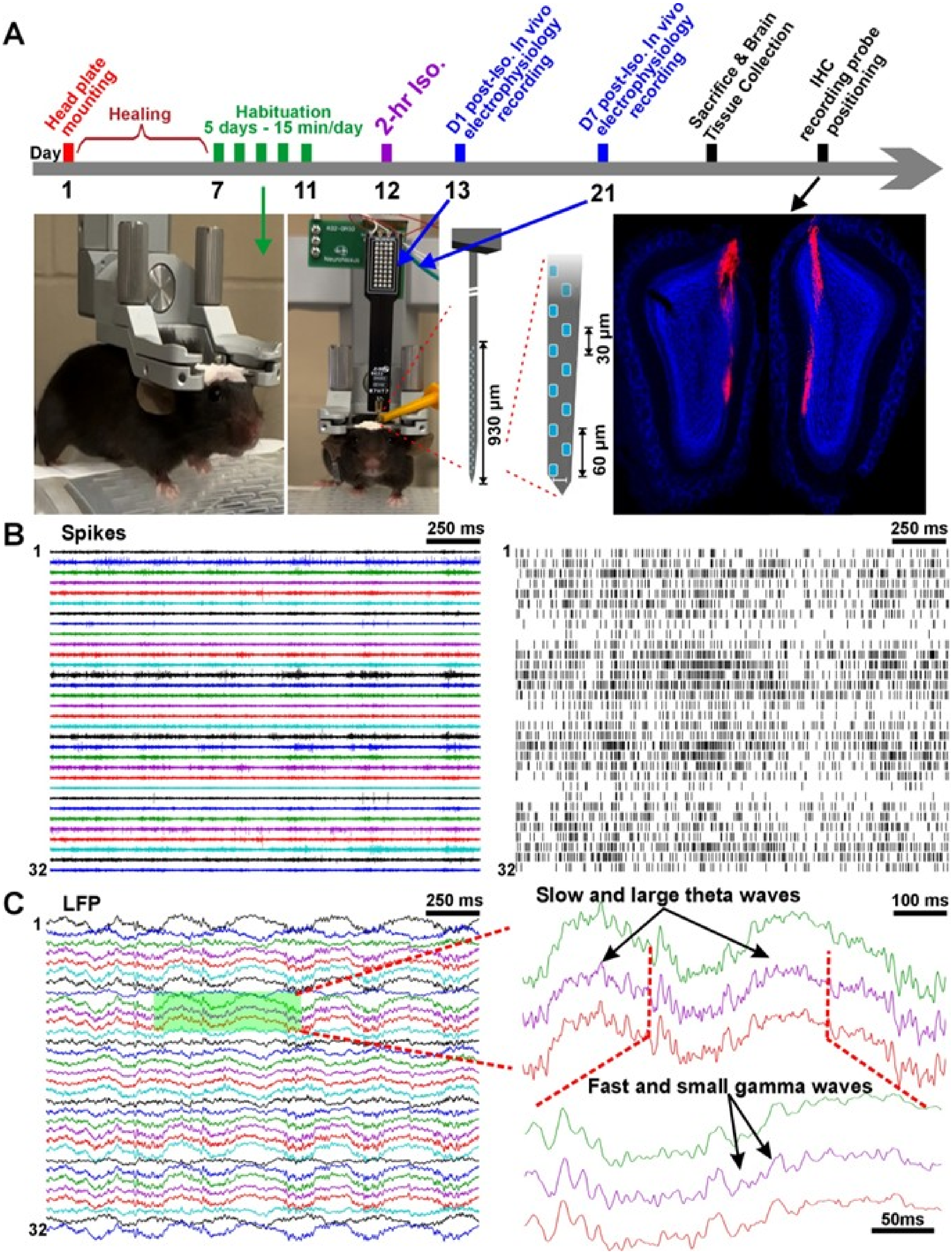
A
Head plate mounting
Healing
Habituation
5 days - 15 min/day
2-hr Iso.
D1 post-Iso. In vivo electrophysiology recording
D7 post-Iso. In vivo electrophysiology recording
Sacrifice & Brain Tissue Collection
IHC recording probe positioning
Day
1
7
11
12
13
21
930 µm
30 µm
60 µm
B
Spikes
250 ms
1
32
250 ms
1
32
C
LFP
250 ms
1
32
Slow and large theta waves
100 ms
Fast and small gamma waves
50ms

**Fig. S5. *In vivo* electrophysiological recordings in awake and head-fixed mice. (A)** schematic illustration of the experimental design. Photos showing recording from an awake and head fixed mouse using 32-channel probe, confocal image verifying the probe penetration path in the OB mitral cell layer. **(B)** typical traces of spike activities (Left) and raster graph showing detected spikes from the left trace (Right). **(C)** typical local field potential (LFP) traces showing the neural oscillations (Left) and blown-up traces showing the detection of slow and large theta (top) or fast and small gamma oscillatory waves (bottom). IHC: Immunohistochemistry, D1 Post-Iso.: Day 1 post-isoflurane, D7 Post-Iso.: Day 7 post-isoflurane.

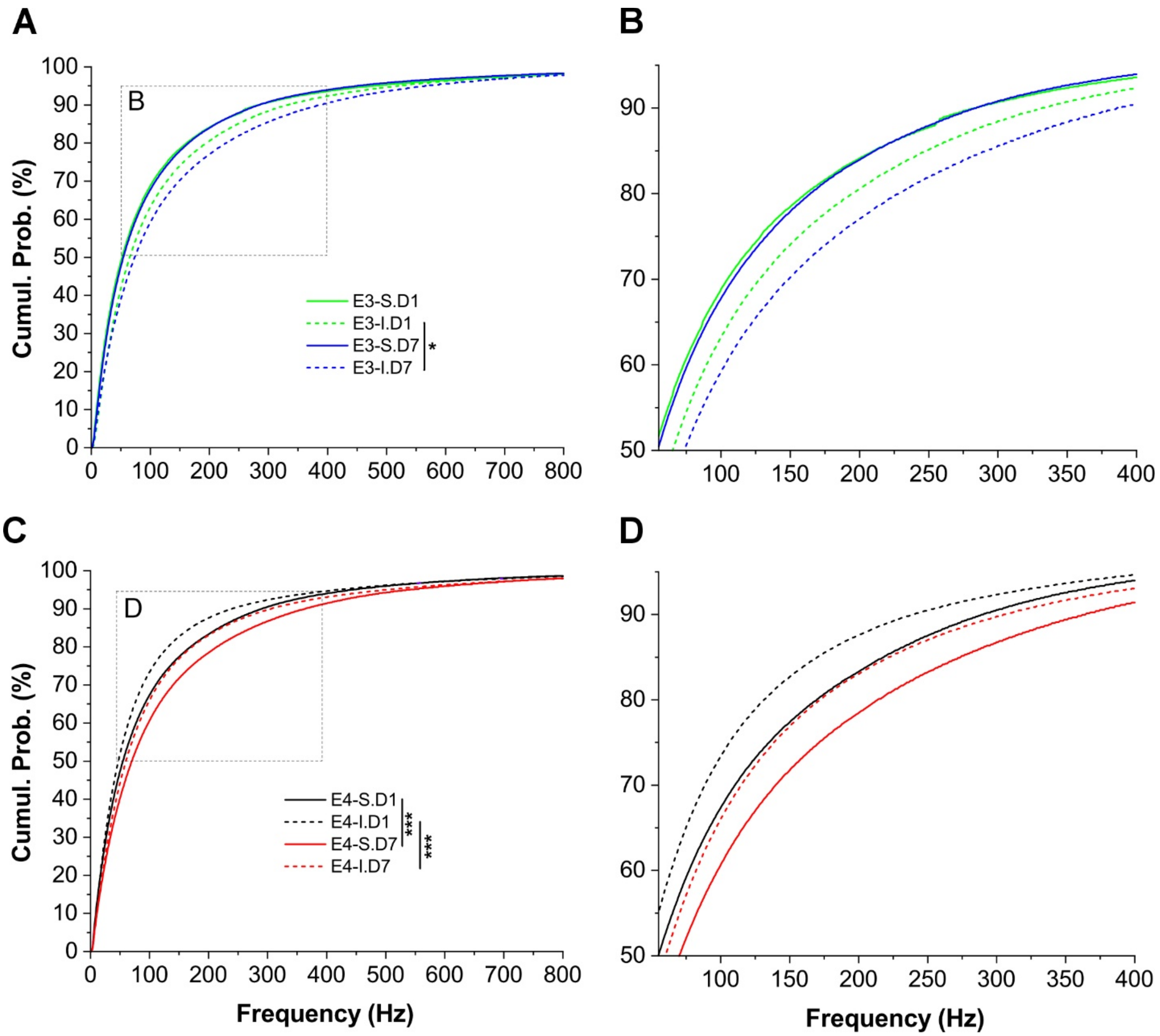


**Fig. S6. Comparison of isoflurane (ISO) effects on neuronal excitability between post-ISO day 1 and day 7. (A)** Comparison of the cumulative probability of spiking frequency in E3 mice of sham (solid green *vs* blue curve) or ISO-exposed (dashed green *vs* blue curve) group between day 1 (D1) and Day 7. **(B)** Blown-up from the dashed box in A. **(C)** Comparison of the cumulative probability of spiking frequency in E4 mice of sham (solid black *vs* red curve) or ISO exposed (dashed black *vs* red curve) group between day 1 (D1) and Day 7. **(D)** Blown-up from the dashed box in C. Comparison between the two groups was analyzed using the Mann-Whitney test. *p<0.05, ***p<0.0001.

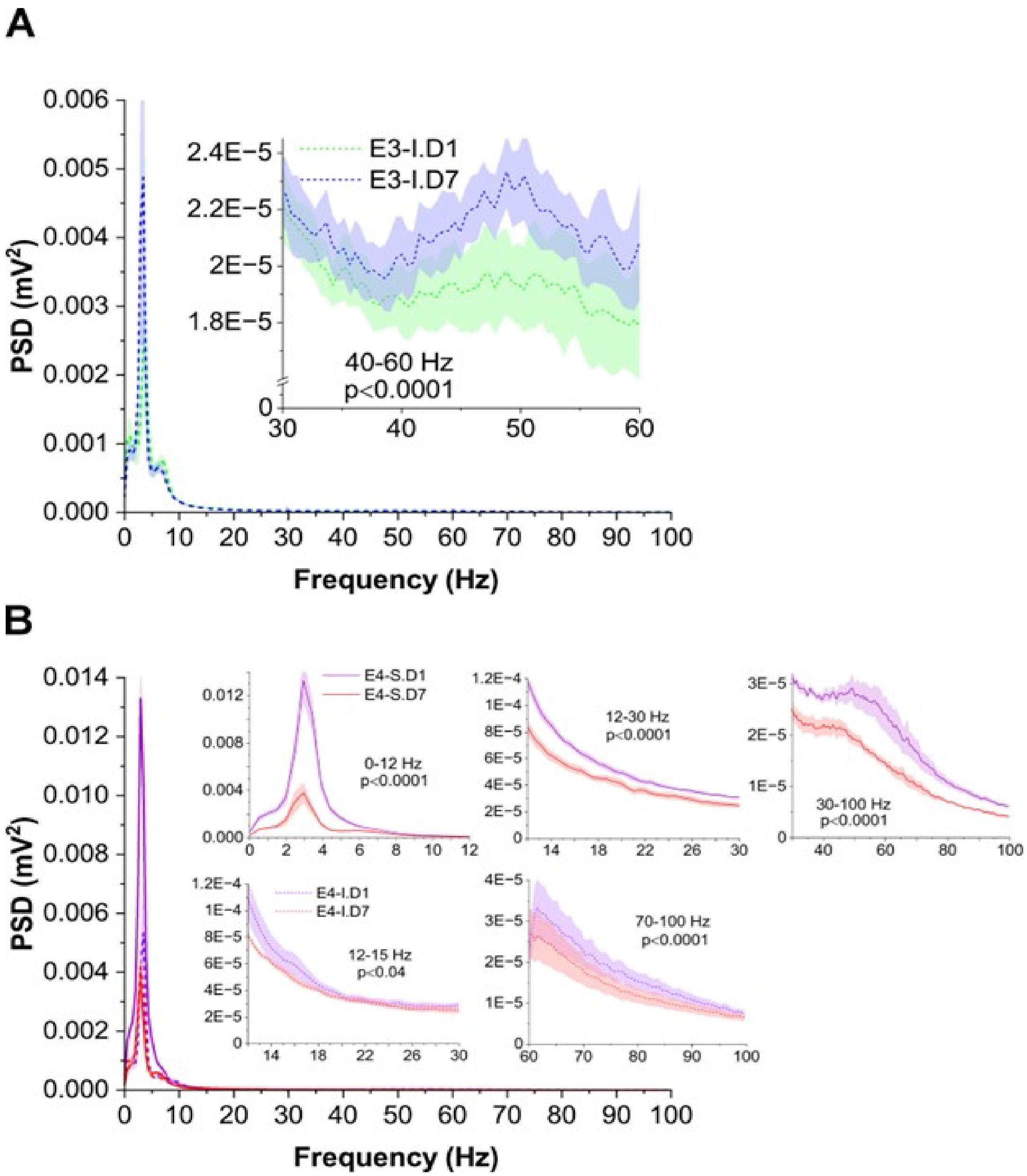


**Fig. S7. Comparison of isoflurane (ISO) effects on neural oscillations of local field potential (LFP) between post-treatment day 1 and day 7. (A)** Comparison of the power spectral density (PSD) of LFP oscillatory activities in E3 mice between day-1 and day-7 post-ISO. The inset is zoom-in from the low gamma (30-60- Hz) frequency band highlighting the difference between two conditions. **(B)** PSD comparison of LFP oscillatory activities between day-1 and day-7 post-ISO in E4 sham or ISO-exposed mice. Insets are zoom-in from each frequency bands δ/□/α (0-12 Hz), β (12-30 Hz), low gamma, (30-60- Hz), and high gamma (60-100 Hz) to highlight differences between day 1 and day 7 groups in sham (solid red vs purple curve, top) or ISO-exposed mice (dashed red vs purple curve, bottom). Each curve (solid or dashed) with shaded area presents mean ± SE. Statistical analysis and comparison were conducted using the Mann-Whitney U test.

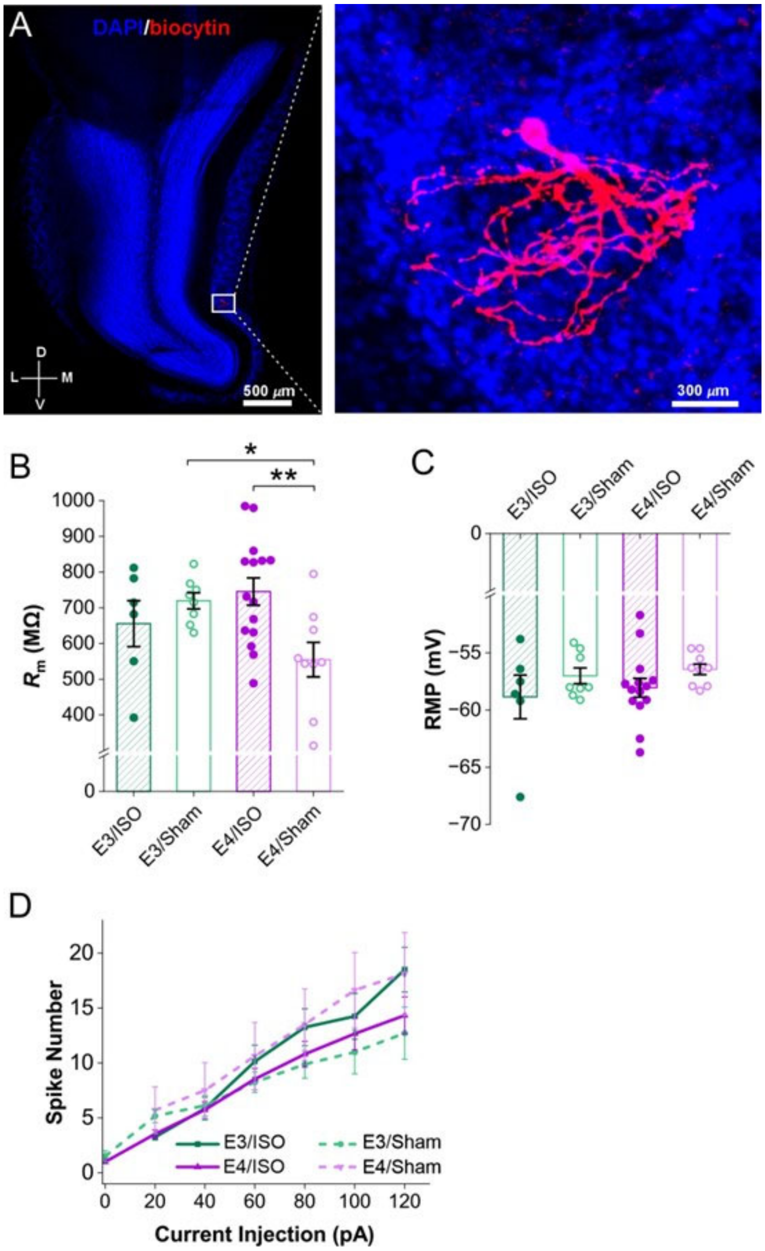


**Fig. S8. Effects on membrane properties of external tufted cells in the olfactory bulb (OB). (A)** Confocal image of a DAPI-counterstained (blue) coronal OB slice with a representative external tufted cell (ETC) filled by biocytin (red) through a patch clamp recording electrode. **(B-D)** comparisons of membrane resistance (Rm, B), resting membrane potential (RMP, C) and number of spikes evoked by injections of stepped depolarizing currents (200 ms, 20 pA/step, D) in ETCs from four groups of mice: E3 with laparotomy and 2-hr isoflurane (E3/ISO), E3 sham (E3/Sham), E4 with laparotomy and 2-h isoflurane (E4/ISO), E4 sham (E4/Sham). Data was presented as mean ± SE and analyzed with Two-way ANOVA and Bonferroni comparisons test. *p<0.05, **p<0.01.

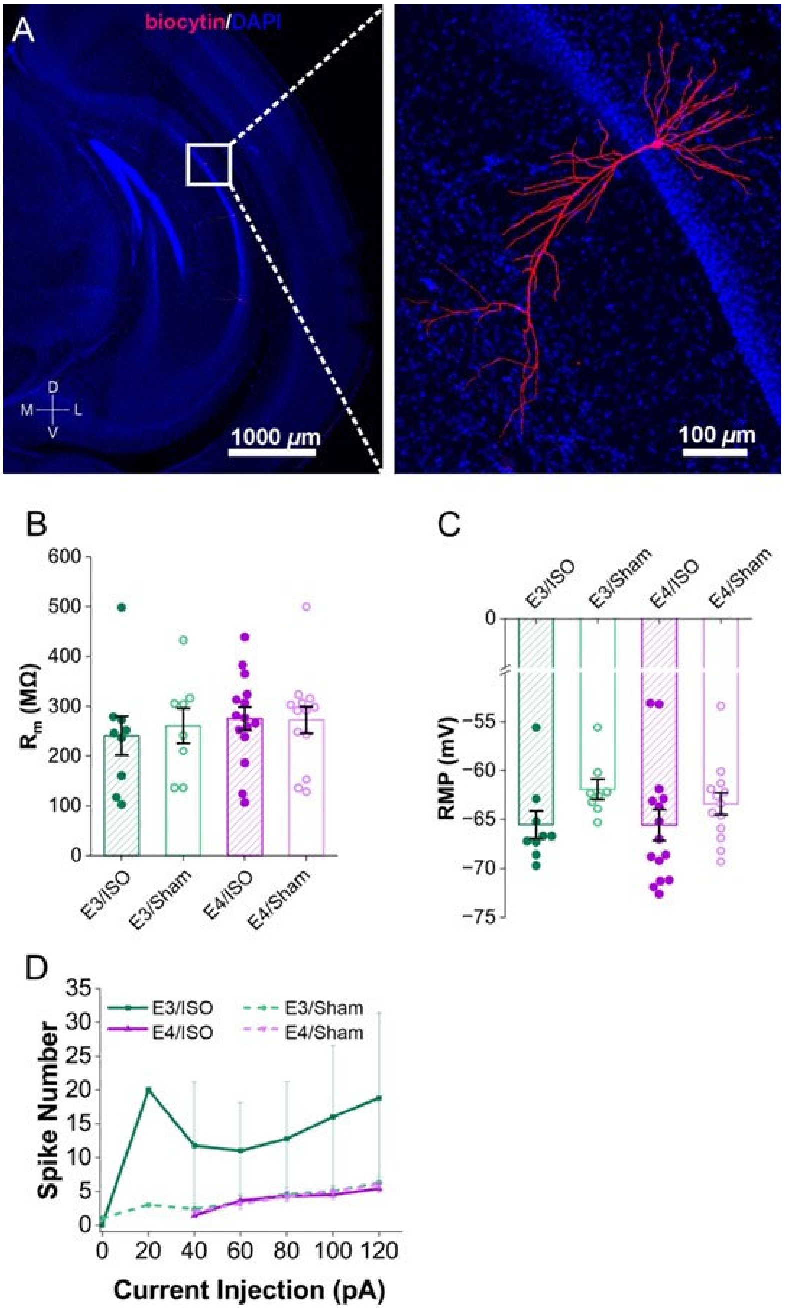


**Fig. S9. Effects on membrane properties of pyramidal neurons in the hippocampus. (A)** Confocal image of a DAPI-counterstained (blue) coronal OB slice with a representative pyramidal neuron filled by biocytin (red) through a patch clamp recording electrode. **(B-D)** Comparisons of membrane resistance (Rm, B), resting membrane potential (RMP, C) and number of spikes evoked by injections of stepped depolarizing currents (200 ms, 20 pA/step, D) in pyramidal neurons from four groups of mice: E3 with laparotomy and 2-hr ISO (E3/ISO), E3 sham (E3/Sham), E4 with laparotomy and 2-h isoflurane (E4/ISO), E4 sham (E4/Sham).

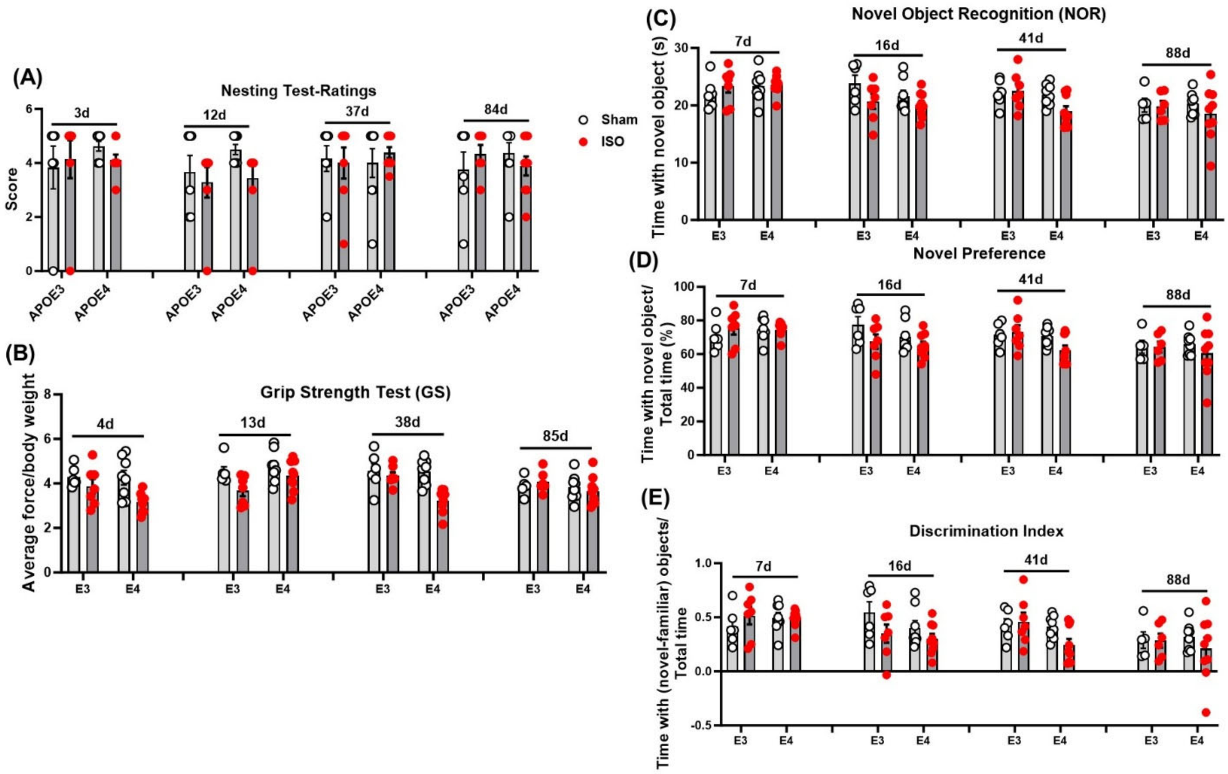


**Fig. S10. Short term isoflurane (ISO) exposure does not affect the general well-being and neuromuscular functions of E4 and E3 mice. (A)** General-welling was examined by the nest-building (NB) test with quantified nest scores. **(B)** Neuromuscular function was unaffected by ISO exposure, as evidenced by grip strength (GS) test. **(C-E)** Non-spatial memory was assessed using the Novel Object Recognition (NOR) test. During the choice phase, the time spent exploring the novel object (C), the animal's preference for the novel object (D), and the discrimination index (E) were recorded. n=6 (E3/Sham), 7 (E3/ISO), 8 (E4/Sham), and 9 (E4/ISO). Data was analyzed with Two-way ANOVA and Tukey's multiple comparisons test.

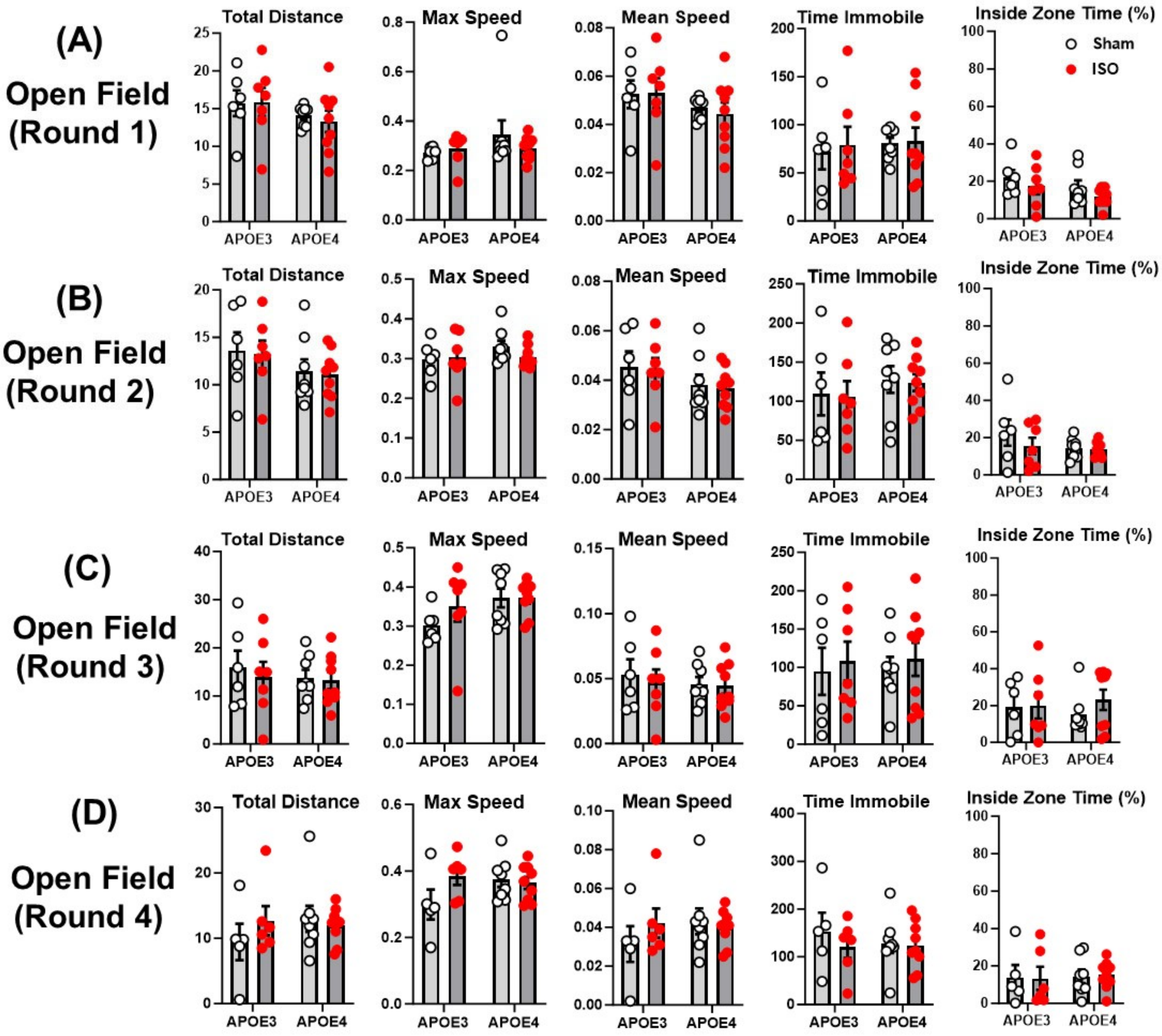


**Fig. S11. Isoflurane (ISO) does not affect the spontaneous locomotor activity of humanized E4 and E3 mice.** The AnyMaze behavior system was used to record and analyze spontaneous activity in an open field apparatus at round 1 **(A)**, round 2 **(B)**, round 3 **(C)** and round 4 **(D)** after ISO exposure. n=6 (E3/Sham), 7 (E3/ISO), 8 (E4/Sham), and 9 (E4/ISO). Data was analyzed with Two-way ANOVA and Tukey's multiple comparisons test.

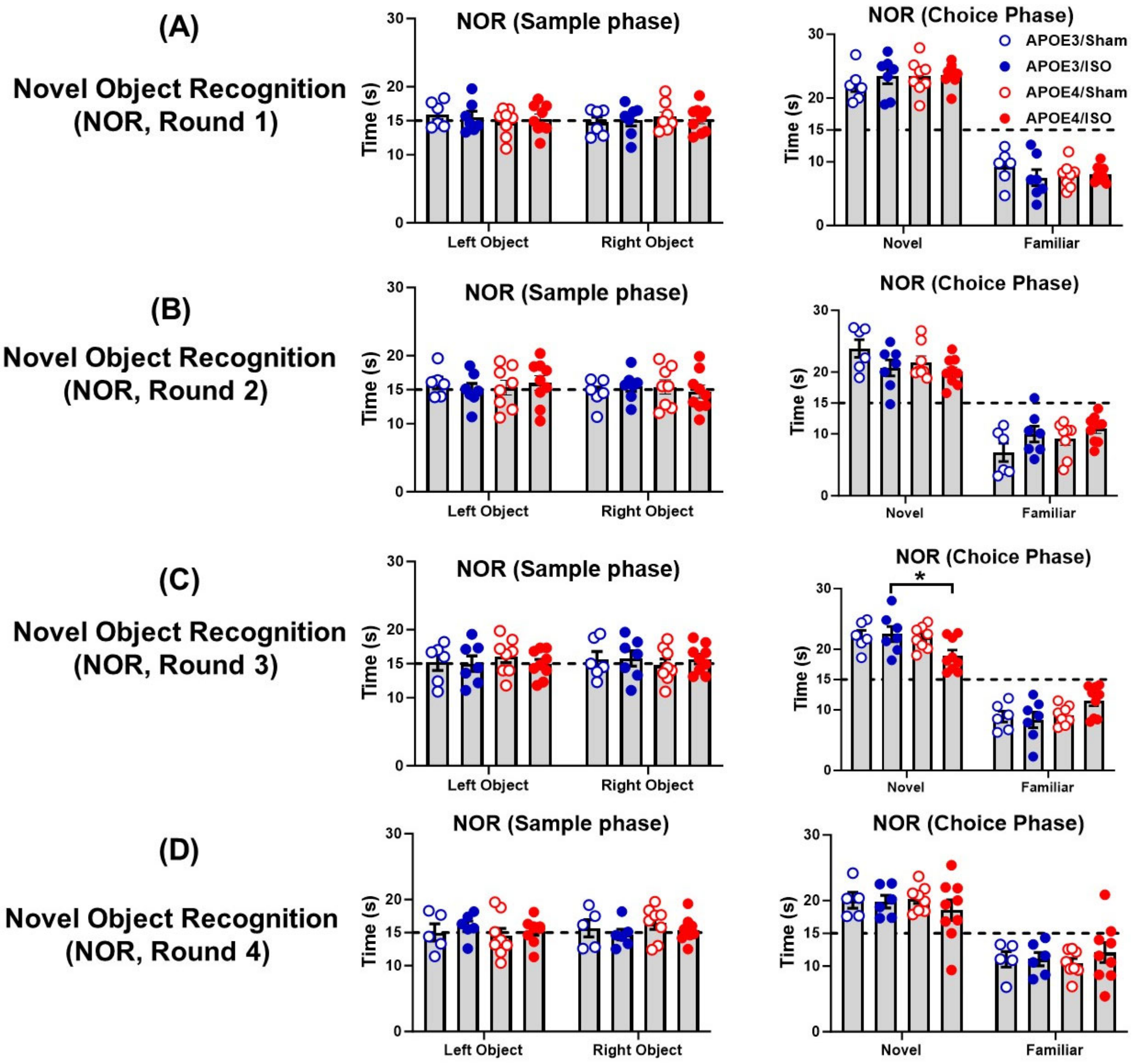


**Fig. S12. There were no differences between the groups in the sample phase of the novel object recognition (NOR) test. (A-D)** A short-term 2h exposure to isoflurane (ISO) with laparotomy did not affect placement preference when exploring the familiar objects but showed a transient impairment on round 3 (C) before recovery on the final round of testing. n=6 (E3/Sham), 7 (E3/ISO), 8 (E4/Sham), and 9 (E4/ISO). Data was analyzed with Two-way ANOVA and Tukey's multiple comparisons test.

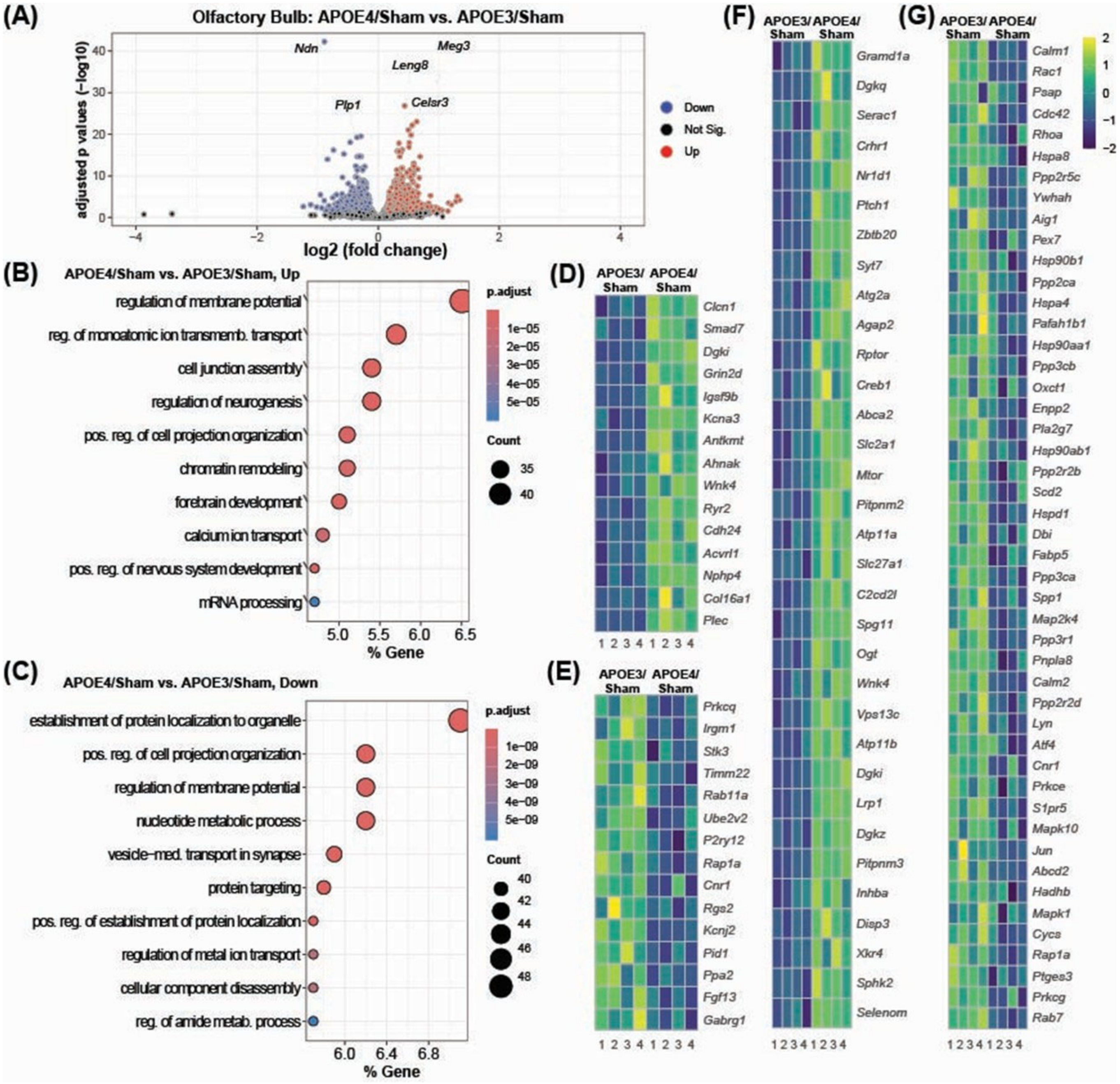


**Fig. S13. Baseline olfactory bulb transcriptomes differ markedly between E4 and E3 mice.** **(A)** Volcano plot displaying differentially expressed genes (DEGs) in the E4/Sham vs. E3/Sham comparison. **(B-C)** Pathway enrichment analysis of up (B) and downregulated (C) DEGs with Gene Ontology molecular processes. **(D-E)** Heatmaps displaying DEGs associated with the top three upregulated pathways (D) and top three downregulated pathways (E) identified between E4/Sham vs. E3/Sham groups. **(F)** Heatmap of genes involved with lipid metabolism. n=4 mice/group.

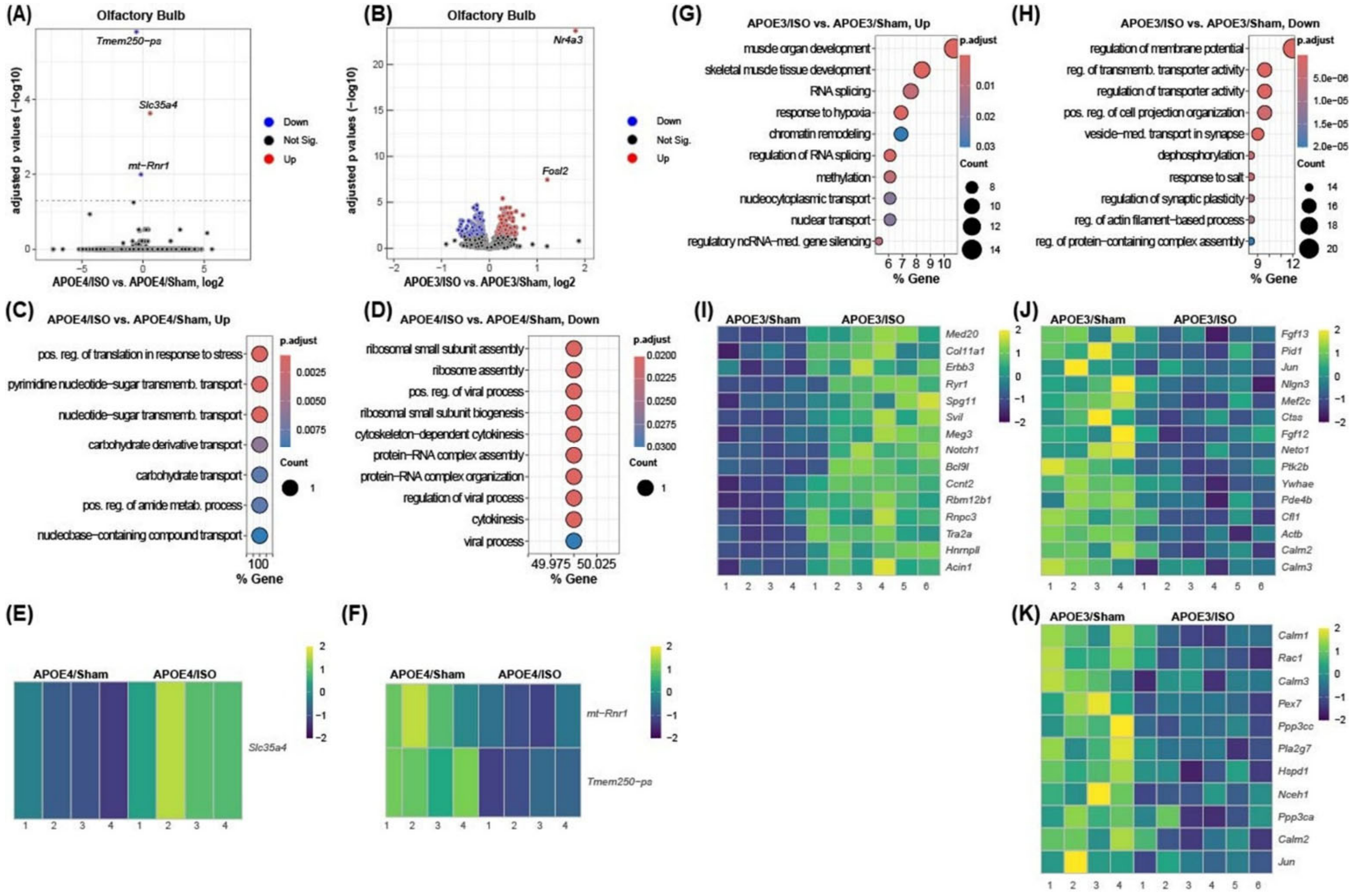


**Fig. S14. RNAseq reveals molecular alterations in the olfactory bulb (OB) 90d after isoflurane (ISO) exposure. (A-B)** Volcano plot of all genes after pairwise comparison of E4/ISO vs. E4/Sham (A) and E3/ISO vs, E3/Sham (B). **(C-F)** Pathway enrichment analysis and heatmap of up- (C, E) and downregulated (D, F) DEGs derived from the E4/ISO vs. E4/Sham comparison. **(G-J)** Pathway enrichment analysis and heatmap of up- (G,I) and downregulated (H, J) DEGs derived from the E3/ISO vs. E3/Sham. n=4 (E4/Sham), 4 (E4/ISO), 4 (E3/Sham), and 6 (E3/ISO).

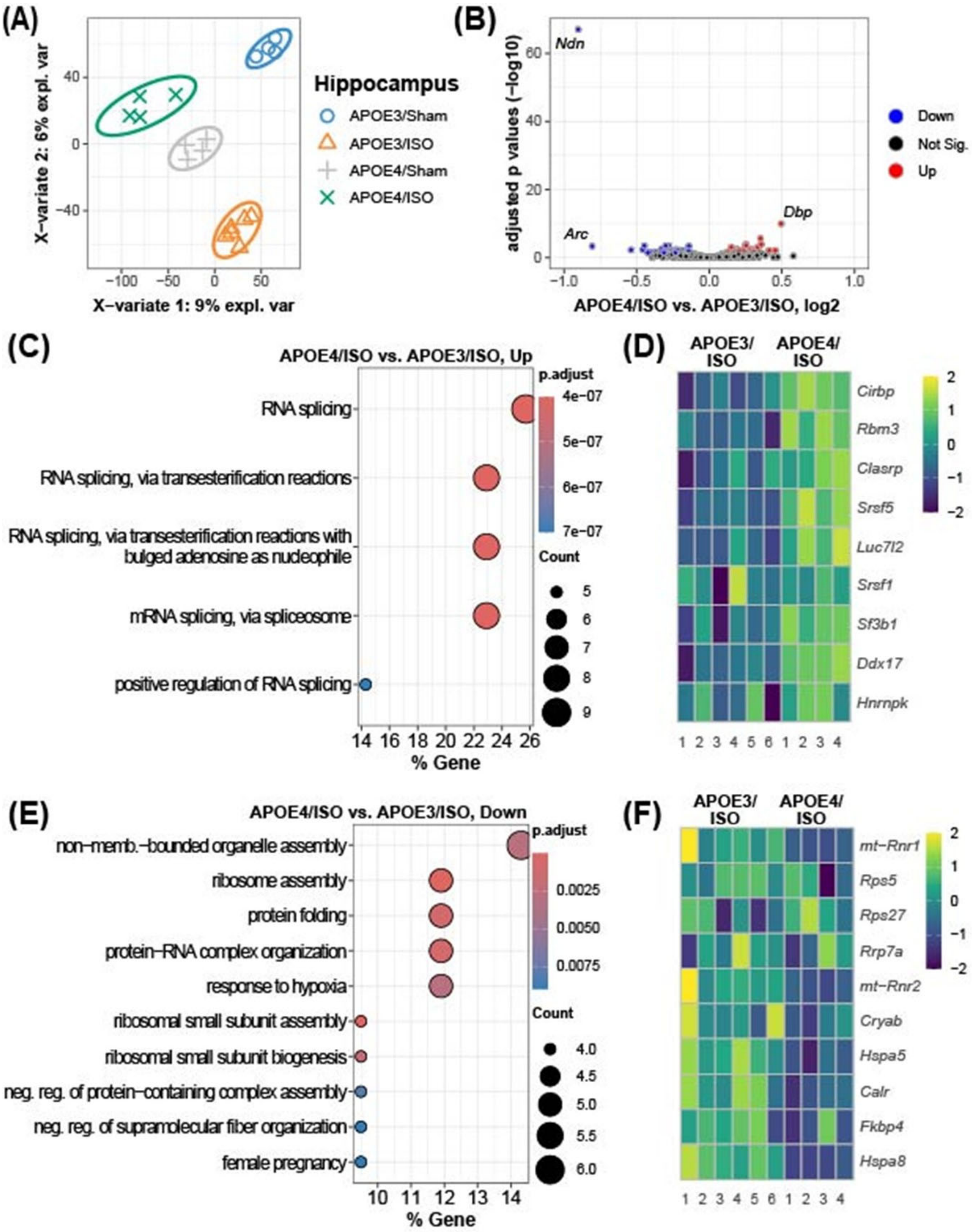
(A)
Hippocampus
APOE3/Sham
APOE3/ISO
APOE4/Sham
APOE4/ISO
X−variate 1: 9% expl. var
X−variate 2: 6% expl. var
(B)
Ndn
Arc
Dbp
Down
Not Sig.
Up
adjusted p values (−log10)
APOE4/ISO vs. APOE3/ISO, log2
(C)
APOE4/ISO vs. APOE3/ISO, Up
RNA splicing
RNA splicing, via transesterification reactions
RNA splicing, via transesterification reactions with bulged adenosine as nucleophile
mRNA splicing, via spliceosome
positive regulation of RNA splicing
p.adjust
Count
% Gene
(D)
APOE3/ ISO
APOE4/ ISO
Cirbp
Rbm3
Clasrp
Srsf5
Luc7l2
Srsf1
Sf3b1
Ddx17
Hnrnpk
(E)
APOE4/ISO vs. APOE3/ISO, Down
non-memb.-bounded organelle assembly
ribosome assembly
protein folding
protein-RNA complex organization
response to hypoxia
ribosomal small subunit assembly
ribosomal small subunit biogenesis
neg. reg. of protein-containing complex assembly
neg. reg. of supramolecular fiber organization
female pregnancy
p.adjust
Count
% Gene
(F)
APOE3/ ISO
APOE4/ ISO
mt-Rnr1
Rps5
Rps27
Rrp7a
mt-Rnr2
Cryab
Hspa5
Calr
Fkbp4
Hspa8

**Fig. S15. E4 mice show distinct transcriptomic signatures in the hippocampus (HI) at 90d following isoflurane (ISO) exposure. (A)** PLS-DA plot of normalized gene expression profiles in E4 and E3 mice under ISO and sham conditions. **(B)** Volcano plot displaying differentially expressed genes (DEGs) in the E4/ISO vs. E3/ISO comparison. **(C-D)** Pathway enrichment analysis of up (C) and downregulated (D) DEGs with Gene Ontology molecular processes. **(E-F)** Heatmaps displaying DEGs associated with the top three upregulated pathways (E) and top three downregulated pathways (F) identified between E4/ISO and E3/ISO groups. Heatmaps illustrate sample-wise variation (columns) across genes (rows). Color intensity reflects relative transcript abundance across samples: warmer colors for higher, cooler for lower. n=4 (E4/ISO) and 6 (E3/ISO) mice.

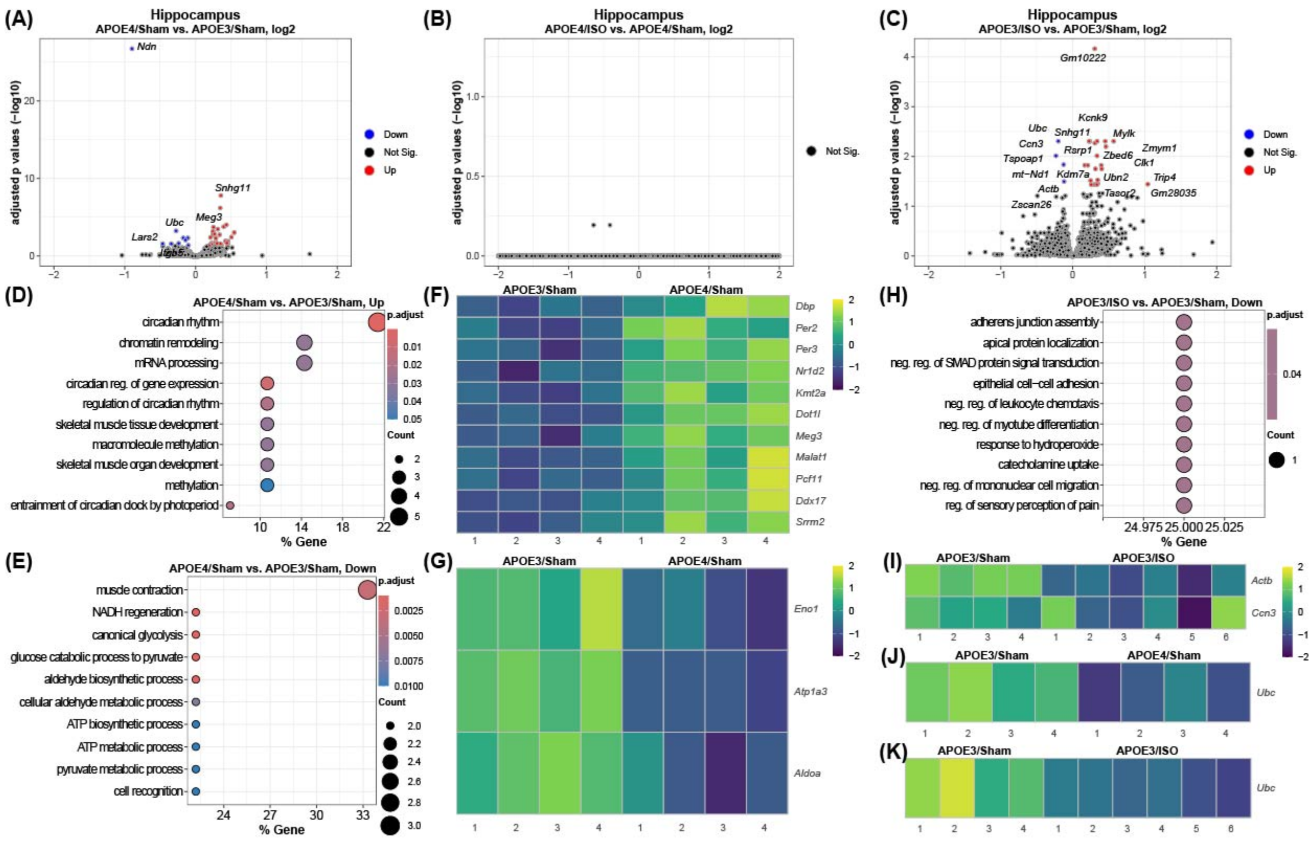


**Fig. S16. Transcriptomic profiling of the hippocampus (HI) reveals distinct baseline and isoflurane (ISO) induced changes in E4 and E3 mice 90 days after exposure.** (A) Volcano plot of all genes after pairwise comparison of E4/Sham vs. E3/Sham (B-E) Pathway enrichment analysis via GO molecular processes of the upregulated (B) and downregulated (D) DEGs and heatmap of genes involved in the top three pathways (C, E). (F) Volcano plot of all genes comparing E4/ISO vs. E4/Sham. (G) Volcano plot of all genes comparing E3/ISO vs. E3/Sham. (H-I) GO molecular process enrichment of downregulated DEGs (H), with corresponding heatmaps of representative genes from the top three pathways (I). n = 4 (E3/Sham), 4 (E4/Sham), 4 (E3/ISO), and 6 (E4/ISO) mice.
